# Sparse matrix factorization robust to sample sharing across GWAS reveals interpretable genetic components

**DOI:** 10.1101/2024.11.12.623313

**Authors:** Ashton R. Omdahl, Joshua S. Weinstock, Rebecca Keener, Surya B. Chhetri, Marios Arvanitis, Alexis Battle

**Affiliations:** Department of Biomedical Engineering, Johns Hopkins University; Baltimore, MD, 21218; Department of Human Genetics, Emory University; Atlanta, GA, 30322, USA; Division of Cardiology, Department of Medicine, Johns Hopkins University, Baltimore, MD, 21205, USA; Department of Computer Science, Johns Hopkins University; Baltimore, MD, 21218, USA; Department of Genetic Medicine, Johns Hopkins University; Baltimore, MD, 21218, USA; Malone Center for Engineering in Healthcare, Johns Hopkins University, Baltimore, MD, 21218, USA; Data Science and AI Institute, Johns Hopkins University, Baltimore, MD, 21218, USA

## Abstract

Complex trait-associated genetic variation is highly pleiotropic. This extensive pleiotropy implies that multi-phenotype analyses are informative for characterizing genetic associations, as they facilitate the discovery of trait-shared and trait-specific variants and pathways (“genetic factors”). Previous efforts have estimated genetic factors using matrix factorization (MF) applied to numerous GWAS. However, existing methods are susceptible to spurious factors arising from residual confounding due to sample-sharing in biobank GWAS. Furthermore, MF approaches have historically estimated dense factors, loaded on most traits and variants, that are challenging to map onto interpretable biological pathways. To address these shortcomings, we introduce “GWAS latent embeddings accounting for noise and regularization” (GLEANR), a MF method for detection of sparse genetic factors from summary statistics. GLEANR accounts for sample sharing between studies and uses regularization to estimate a data-driven number of interpretable factors. GLEANR is robust to confounding induced by shared samples and improves the replication of genetic factors derived from distinct biobanks. We used GLEANR to evaluate 137 diverse GWAS from the UK Biobank, identifying 58 factors that decompose the genetic architecture of input traits and have distinct signatures of negative selection and degrees of polygenicity. These sparse factors can be interpreted with respect to disease, cell-type, and pathway enrichment. We highlight three such factors capturing platelet measure phenotypes and enriched for disease-relevant markers corresponding to distinct stages of platelet differentiation. Overall, GLEANR is a powerful tool for discovering both trait-specific and trait-shared pathways underlying complex traits from GWAS summary statistics.

## Introduction

Genome-wide association studies (GWAS) have identified thousands of associations between common genetic variants and complex traits^1,2^. Efforts to understand genetic relationships between diseases often leverage pairwise comparison of GWAS^3,4^ and have found evidence of widespread pleiotropy among genetic loci^5,6^. Studies of pleiotropy focus either on individual variants^4,5^ or examine the genetic correlation^3^ between GWAS, which detects covariance between the genetic effects of two phenotypes. Genetic correlation may arise from common biology among traits at multiple scales, such as shared variants, pathways, and cell types ^7,8^. Pleiotropy can thus serve to support identification of important genetic pathways with multi-trait impact. Several studies^7–13^ have shown that these relationships can be captured by latent genetic components (factors). These factors map genetically related traits to groups of variants belonging to shared pathways or cell types potentially missed by genome-wide analysis or narrow consideration of single variants.

Most study designs for estimating such genetic factors across many traits employ matrix factorization (MF) approaches ^7,8,10,13^. MF explicitly decomposes an input matrix of GWAS effects for all traits into the product of two matrices: the first matrix maps SNPs to latent factors, and the second matrix maps latent factors to traits. While MF approaches are accessible and scalable, existing studies have a few notable limitations. First, standard approaches yield dense factors, where every trait is loaded on every factor, which in turn has non-zero effects for every SNP. This presents a challenge for interpretation, as the factors are less likely to represent distinct pathways, cell-types, or subsets of affected traits, and may increase the susceptibility to overfitting. Second, present studies have not consistently accounted for sources of potential confounding in GWAS data, including non-independence of target studies due to sharing of samples. Finally, downstream analysis of factors has focused primarily on characterization of the top variants and traits for each factor, while properties such as factor genetic architecture or distinct contributions of trait-relevant cell types remain unexplored.

To estimate interpretable genetic factors that are robust to shared samples between studies, we developed GWAS latent embeddings accounting for noise and regularization (GLEANR). GLEANR uses dynamic model selection to yield sparse factors while statistically accounting for confounding covariance from sample sharing and variation in estimation error. In simulations, GLEANR outperforms other factorization methods when input studies share samples. By adjusting for the effects of sample sharing, our approach successfully estimates genetic components which are more replicable across distinct cohorts. In an evaluation of 137 GWAS traits across the phenome from the UK Biobank^14,15^ (UKBB), GLEANR identifies genetic factors with distinct tissue enrichments, polygenic architectures, and signatures of selection. Among these, we identify three factors related to platelet traits which capture distinct stages of platelet formation related to inherited platelet disorders. GLEANR is a powerful tool for hypothesis-free exploration of genetic variants impacting shared pathways among component traits.

## Results

### Overview of GLEANR model and pipeline

Complex traits are highly pleiotropic^3–6,16,17^. To capture common genetic pathways or components among multiple traits, we developed GLEANR, an MF approach tailored to GWAS summary statistics. GLEANR estimates sparse latent genetic components, decomposing the full set of genetic effects across GWAS into a lower dimensional representation consisting of factors shared by traits. This procedure accounts for both the uncertainty inherent in GWAS effect size estimates and the sharing of samples between studies.

GLEANR decomposes an input matrix of LD-pruned marginal effect sizes of *N* SNPs on *M* GWAS traits, **B** ∈ℝ^*N*×*M*^, as

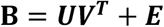

where ***V*** ∈ℝ^*M*×*K*^ weights phenotypes on *K* genetic factors and ***U*** ∈ℝ^*N*×*K*^ reflects the SNP effects contributing to those factors. Both ***U*** and ***V*** are sparse, as not all phenotypes are genetically correlated, nor do all SNPs make pleiotropic contributions to these shared effects^4^. Hence, these matrices are estimated with sparsity-inducing L_1_ regularization^18,19^. However, to account for pleiotropic genetic associations that impact many traits, the first factor is not subject to regularization in ***V***, and thus weighted on all traits (a ubiquitous factor). Each row of residual noise matrix ***E*** (***ϵ***_**1**_, ***ϵ***_**2**_, … ***ϵ***_***N***_) corresponds to a single SNP across all studies and reflects both measurement uncertainty and non-genetic correlation between studies (e.g. sharing of samples). This is modelled as ***ϵ***_***n***_**∼***MVN*_*M*_ (0, diag(***s***_***n***_) × ***C*** × diag(***s***_***n***_)=, where 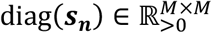 is a diagonal matrix of GWAS standard errors for SNP *n* and ***C*** is a block matrix of correlation induced by sample sharing between studies, which may be estimated directly from summary statistics using cross-trait LD-score regression^3^ (XT-LDSR; Fig. 1A).

**Figure 1:**
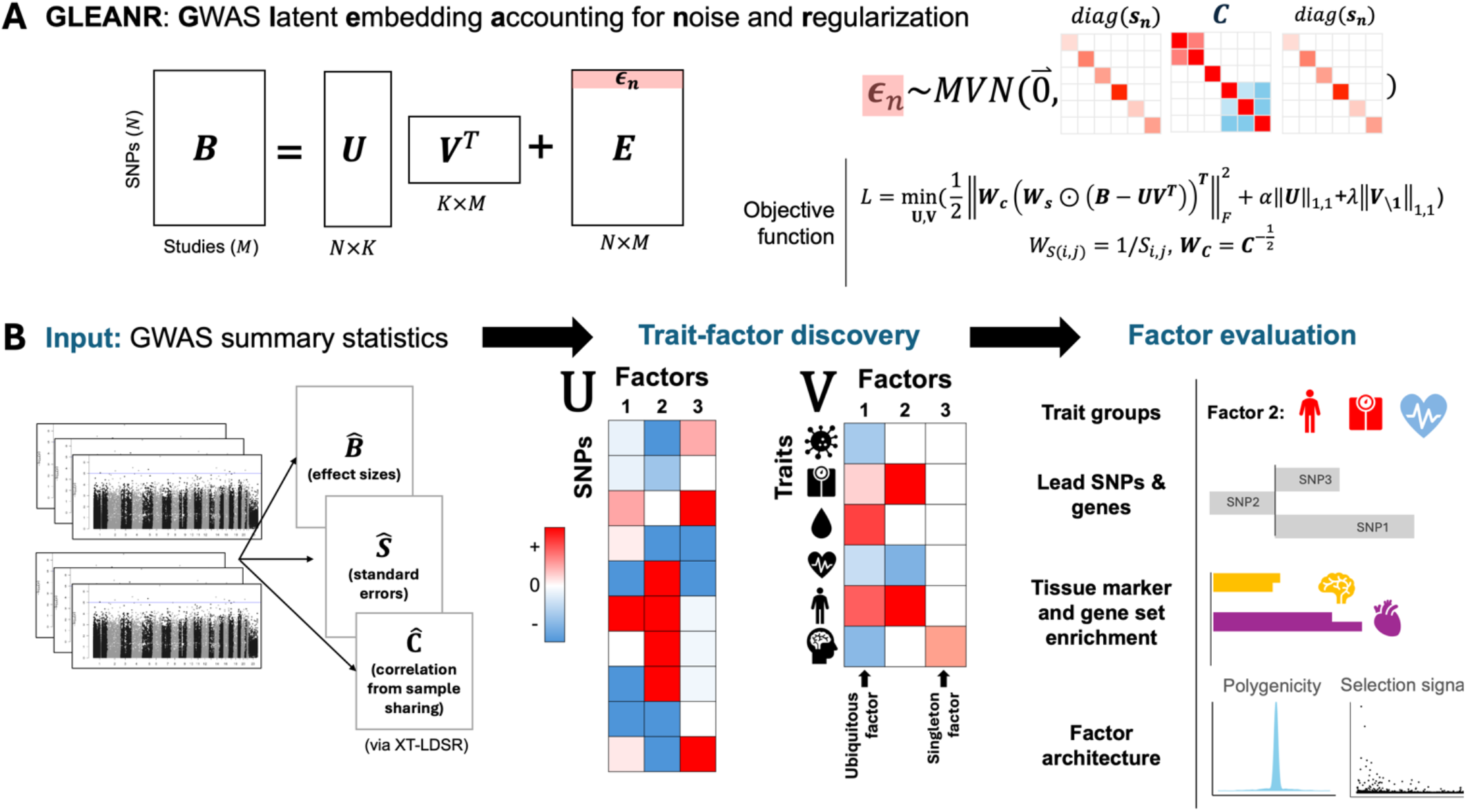
GLEANR decomposes GWAS summary statistics into sparse latent genetic factors. **A**) GLEANR maps GWAS SNPs (in ***U***) and studies (in ***V***) onto a latent *K*-dimensional space. Covariance of a single SNP *n* across studies is modeled as the matrix product of SNP standard errors (*diag*(***s***_***n***_)) and a matrix of correlations induced by study sample sharing (***C***). GLEANR minimizes the indicated objective function, weighting terms by the inverse of standard errors (***W***_***s***_) and the decorrelating transformation ***W***_***c***_ derived from ***C***. B) GWAS summary statistics provide estimates of SNP effect sizes 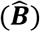 and standard errors 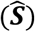 and are used to estimate correlation due to sample sharing 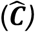. GLEANR performs model selection and fitting to select *K* factors loaded on traits and SNPs. A latent factor consists of traits (columns of ***V***) and loaded SNPs (the corresponding columns of ***U***). The first factor spans all traits (ubiquitous factor) and some factors may load on only a single trait (singleton factors). We evaluate factors by considering their member traits, top SNPs, genes, tissue enrichments, and factor architecture.

To estimate ***U*** and ***V***, GLEANR minimizes the bi-convex objective function in Fig. 1A through an iterative procedure of alternating generalized least squares regression. One of ***U*** or ***V*** is fixed to estimate the other via regression against **B**, with matrix elements weighted by standard errors (***W***_***s***_) and transformed by a decorrelating matrix (***W***_***c***_) derived from ***C***. This weighting and decorrelation help eliminate uninformative or spurious factors. GLEANR dynamically selects sparsity weights (*α* and *λ*) by minimizing the Bayesian Information Criteria (BIC) during alternating regression steps^20–22^.

Factors can be directly interpreted by examining the non-zero entries in ***U*** and ***V***, along with their magnitude and direction, or may be used in downstream analysis as latent representations of genetic pathways shared by traits. Factors may capture narrow trait-specific contribution of variants, such as in singleton factors loaded on just one study, or may reflect broad variant effects, as in the unregularized ubiquitous factor (Fig. 1B). As representations of genetic pathways, factor SNP weights can capture distinct genetic architectures with tissue-marker and functional enrichments.

### Modest cohort overlap results in spurious factors for standard MF methods

Overlap among samples between GWAS is commonplace. Existing MF studies of biobank summary statististics^10,23^ commonly include GWAS with substantial sample sharing, as reported by estimates on the UKBB Genetic Correlation Portal^3,24^ (Web Resources; Fig. S1). To characterize the impact of sample sharing between GWAS on MF, we evaluated several factorization and model selection methods on GWAS derived from simulated, correlated phenotypes with no causal genetic component. By omitting a genetic signal, we isolated the impact of sample sharing since any discovered factors would be spurious. We evaluated methods including the singular value decomposition (SVD), testing both the elbow point of eigenvalues (SVD (elbow)) and factors corresponding to eigenvalues greater than the average eigenvalue^10^ (SVD (avg)) to select the number of factors (K). We also added our own modification of the SVD in which we adjusted input Z-scores with a decorrelating transformation^4,25^ to account for sample sharing (SVD-adj), as a baseline method that does adjust for sample sharing. Finally, we included the empirical Bayesian method flash^26^ which automatically performs model selection. We compared these methods with GLEANR, both with covariance adjustment (standard GLEANR) and with no adjustment (unadjusted GLEANR or GLEANR-U). In these simulations, GLEANR and SVD-adj were provided with a cohort overlap correlation matrix estimated from phenotype correlations and the number of overlapping samples^25^ (Methods).

We simulated genotypes based on minor allele frequencies (MAF > 0.01) reported for Europeans in the UKBB^14,27^ among LD-pruned HapMap3^28^ SNPs. For phenotypes, we randomly simulated values from a Gaussian distribution with mean and variance based on estimates from European UKBB^15^ individuals for height. We simulated two additional related phenotypes roughly based on the relationship between height, weight, and body mass index (BMI), assuming weight is linearly related to height and BMI is based on their ratio, both with added Gaussian noise (Methods). This approach ensured that correlations between simulated phenotypes were reasonably approximate to those in real traits. We then performed linear regression between the simulated phenotypes and genotypes, estimating summary statistics in three distinct populations for a total of nine GWAS, each conducted across 30,000 simulated individuals at increasing levels of sample sharing.

Given the independent generation of genotypes and phenotypes, no true genetic factors influence traits in this simulated data. In this scenario, we found that sharing of less than 25% of study individuals was sufficient to cause baseline MF model selection procedures to nominate spurious components (Fig. 2A). By contrast, the covariance adjustment procedure used in GLEANR avoided detection of spurious factors even at high levels (>90%) of shared samples (Fig. 2A, dark orange, far right panel). While GLEANR-U also avoided detection of spurious factors better than most other methods (light orange) at low and moderate sample overlap, presumably due to enforced sparsity, at the highest levels of sample overlap GLEANR-U also yielded spurious factors. Both sparsity-inducing penalty and direct adjustment contribute to GLEANR performance in the presence of cohort overlap. Thus, in the absence of genetic signal, sample sharing results in the detection of spurious factors unless appropriate statistical adjustment is made.

**Figure 2:**
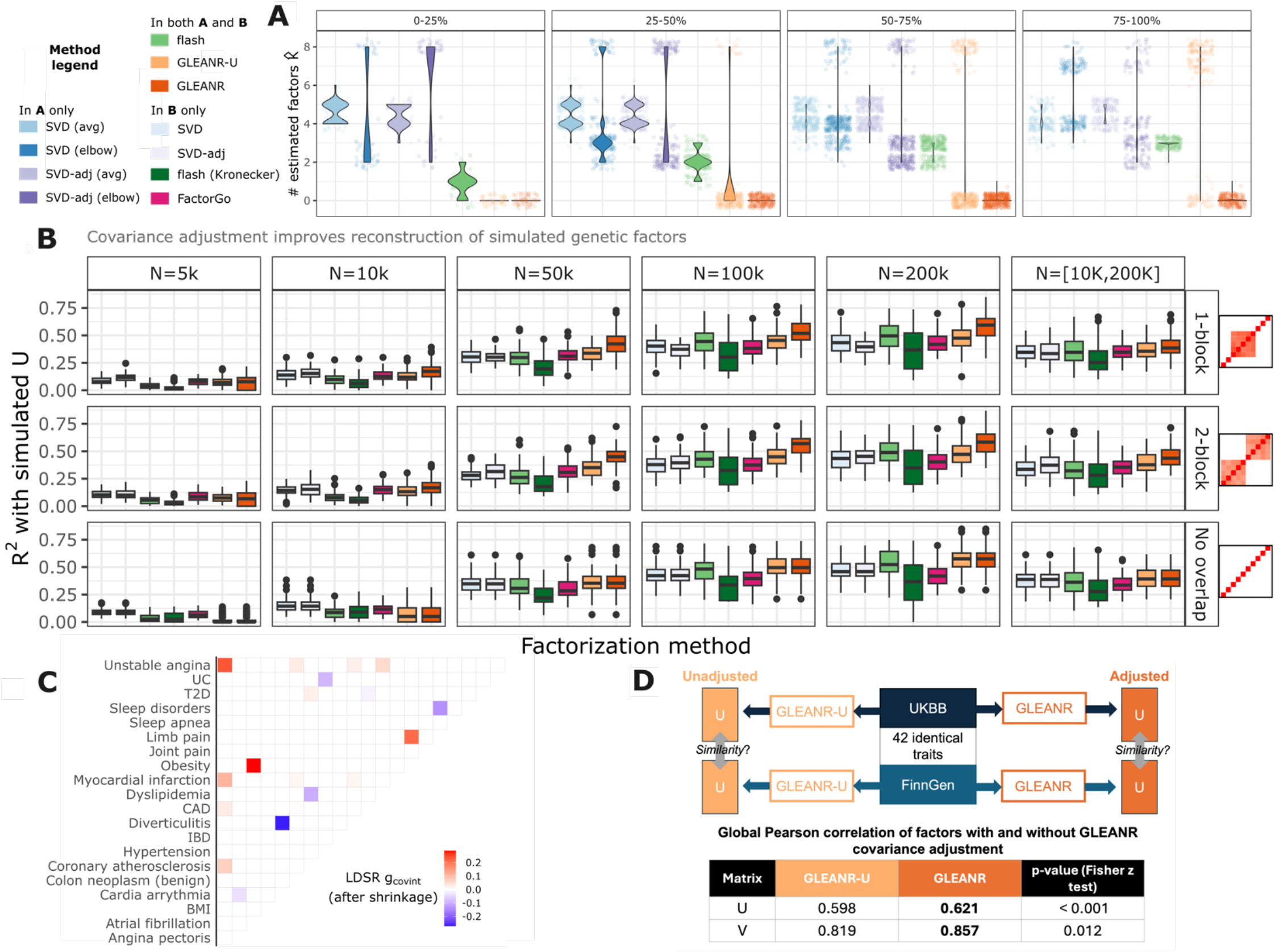
GLEANR more accurately characterizes latent genetic factors in simulation than other tested methods and harmonizes genetic components across cohorts. A) Violin plot of the number of selected factors from various matrix factorization (MF) methods evaluating simulated GWAS with no genetic component at increasing levels of sample sharing between studies. Each point represents a single MF from one of 840 simulations. Facets along the top indicate the prevalence of sample sharing, with 100% indicating that the same samples were used to simulate all phenotypes within a given cohort. B) Variance explained (**R**^2^) by estimated 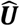 with respect to a known ***U*** (y-axis) across simulations, at varying sample sizes (facet labels at the top), different covariance block structures (facets on the right), across different factorization methods (x-axis, boxplot color). Summary statistics were simulated from sparse ***U*** and ***V*** matrices. Larger **R**^2^ indicates better reconstruction of ***U***. C) Upper triangle heatmap of differences in Finngen and UKBB estimates of correlation due to GWAS sample sharing. Only those with non-zero differences (following variance scaling and GLEANR covariance shrinkage) are shown here. D) Visual overview of analysis comparing UKBB versus FinnGen cohorts. Table reports global Pearson correlation between ***U*** from each cohort, estimated using GLEANR with (dark orange) or without (light orange) covariance adjustment. The GLEANR covariance adjustment procedure yields a more similar genetic basis across cohorts. (Trait abbreviations: BMI, body mass index; CAD, coronary artery disease; IBD: irritable bowel disease; T2D, type 2 diabetes; UC, ulcerative colitis)

### GLEANR recapitulates ground truth genetic factors even with GWAS sample sharing

To evaluate MF estimation of known genetic factors, we simulated GWAS summary statistics from prespecified ***U*** and ***V*** as ***B*** = ***UV***^*T*^ + ***E***. Our simulated ***U*** and ***V*** consisted of five factors drawn from a sparse genetic architecture (Methods). We added to this noise (***E***) drawn from a multivariate normal distribution, with covariance reflecting simulated standard error across multiple sample sizes (from 5,000 to 200,000) and one of three cohort overlap structures (single block, two block, and no overlap, far right of Fig. 2B). These overlap structures correspond to estimates of sample sharing found in UKBB GWAS factorization studies^10,23^ (Fig. S1A). In addition to the SVD and SVD-adj, we evaluated flash with both the default and the “Kronecker” covariance setting, which accounts for more complex residual covariance structures, and FactorGo^23^, a variational model for large-scale GWAS factorization. GLEANR and SVD-adj were provided with the simulated cohort overlap correlation matrix. To assess fit, we matched columns of 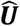 and 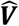 to those in ***U*** and ***V*** by maximizing the correlation between factor pairs, and then calculated the R^2^ across all matched factors after scaling each to unit norm.

In simulations with no cohort overlap (Fig. 2B, bottom row), GLEANR performed comparably to other methods on GWAS with over 10,000 samples. In simulations with either single-block or two-block sample overlap correlation, GLEANR reconstructions of ***U*** (Fig. 2B) had higher median R^2^ than all tested MF methods in sample sizes above 10,000, with similar performance for ***V*** (Fig. S2A). This was consistent with other metrics of fit, including Cohen’s Kappa, which we used to measure discrimination between signed and zero matrix elements (Figs. S2B and S2C). Simply decorrelating Z-scores (SVD-adj) did not consistently improve reconstruction of the latent space relative to unadjusted SVD. At sample sizes below 10,000, GLEANR was slightly outperformed by SVD-based approaches. This was due to sparsity constraints removing signals for small or noisy effects, suggesting that GLEANR may not perform better than baseline methods for GWAS with very small sample sizes. Overall, GLEANR is robust to simulated confounding due to sample sharing between studies and yields improved reconstruction of genetic factors relative to other common methods.

### Sample sharing adjustment improves concordance of factors between distinct cohorts

To evaluate GLEANR ‘s covariance adjustment procedure on published GWAS, we assessed the replication of factors discovered from multiple traits in two distinct cohorts: UKBB ^27^ and FinnGen ^29^. Specifically, the two large cohorts have no samples in common, but within each cohort multiple traits are measured over highly overlapping sets of individuals. We hypothesized that adjusting for sample sharing within each study during factorization would yield genetic factors that were more consistent between cohorts, reflecting more accurate factors due to elimination of confounding from sample overlap.

We selected 42 diverse phenotypes with summary statistics publicly available from both the UKBB^14,24,30^ and FinnGen^29^ to evaluate with GLEANR (Table S1). Traits were chosen to have strong genetic correlation (*r*_*g*_ ≥ 0.8, with adjusted *p* < 0.01) between cohorts. Estimates of sample sharing were generated using XT-LDSR^3^, and adjusted to account for uncertainty and trait heritability (see Methods). These estimates reflected distinct cohort-specific sample sharing structure (differences visualized in Fig 2C), especially among several cardiac phenotypes (e.g. angina, myocardial infarction, CAD). We applied GLEANR and GLEANR-U to GWAS across traits from FinnGen, and then separately to GWAS from the UKBB. To minimize differences related to model selection, we initialized GLEANR at *K*_*init*_ = 41 for both cohorts.

To determine how correcting for cohort-specific sample overlap affected detection of genetic factors, we compared the estimated factors (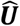 and 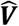) and reconstructed effect sizes 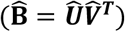 between UKBB and FinnGen. GLEANR nominated both fewer total and a more consistent number of final factors than GLEANR-U (13/12 vs 19/17) and yielded a 4% reduction in overall RRMSE between UKBB and FinnGen reconstructed effect sizes. Globally, GLEANR factors were more strongly correlated across cohorts than GLEANR-U factors (0.62 vs 0.59 for ***U***, Fisher’s Z-test p < 0.001, Fig 2D), scaled to unit length. This improvement in factor similarity is consistent with the estimated magnitude of structural differences in cohort overlap between UKBB and FinnGen (Fig. 2C, Fig. S3). Our results suggest that GLEANR factors are more accurate and less reflective of confounding, non-genetic effects.

GLEANR removed multiple singleton factors from both UKBB and FinnGen (8 and 5, respectively) unadjusted analyses, including a factor loaded on lipo-protein metabolism disorders present in both cohorts (Fig. S4). Importantly, a key heart disease related factor connecting angina pectoris, heart attack, ischemic heart diseases and coronary atherosclerosis remained across cohorts even after covariance adjustment, despite also being influenced by sample sharing. This demonstrates that GLEANR is able to preserve pleiotropic signals between traits while still adjusting for cohort-specific patterns of sample sharing.

### GLEANR across the human phenome identifies 57 sparse factors and one ubiquitous factor

To explore distinct genetic factors spanning the human phenome, we evaluated 137 diverse GWAS conducted on Europeans from the UKBB^14,27^, chosen so none had pairwise genetic correlation *r*_*g*_ > 0.7 (see Methods). These traits span 19 trait categories^31^ (Fig 3A), including metabolic, skeletal, behavioral, respiratory, and immunological phenotypes (Table S2). Across these, we evaluated HapMap3^28^ SNPs with MAF > 0.01 that achieved a nominal *p* < 1 × 10^−4^ in at least one study, which we then LD clumped resulting in a total of 41,259 variants. Our LD clumping procedure prioritized inclusion of variants achieving nominal significance in the largest number of target studies. Overall, only 1,455 variants achieve this p-value threshold in 10 or more traits, with nearly half of SNPs (20,137) passing this threshold in just one trait.

**Figure 3:**
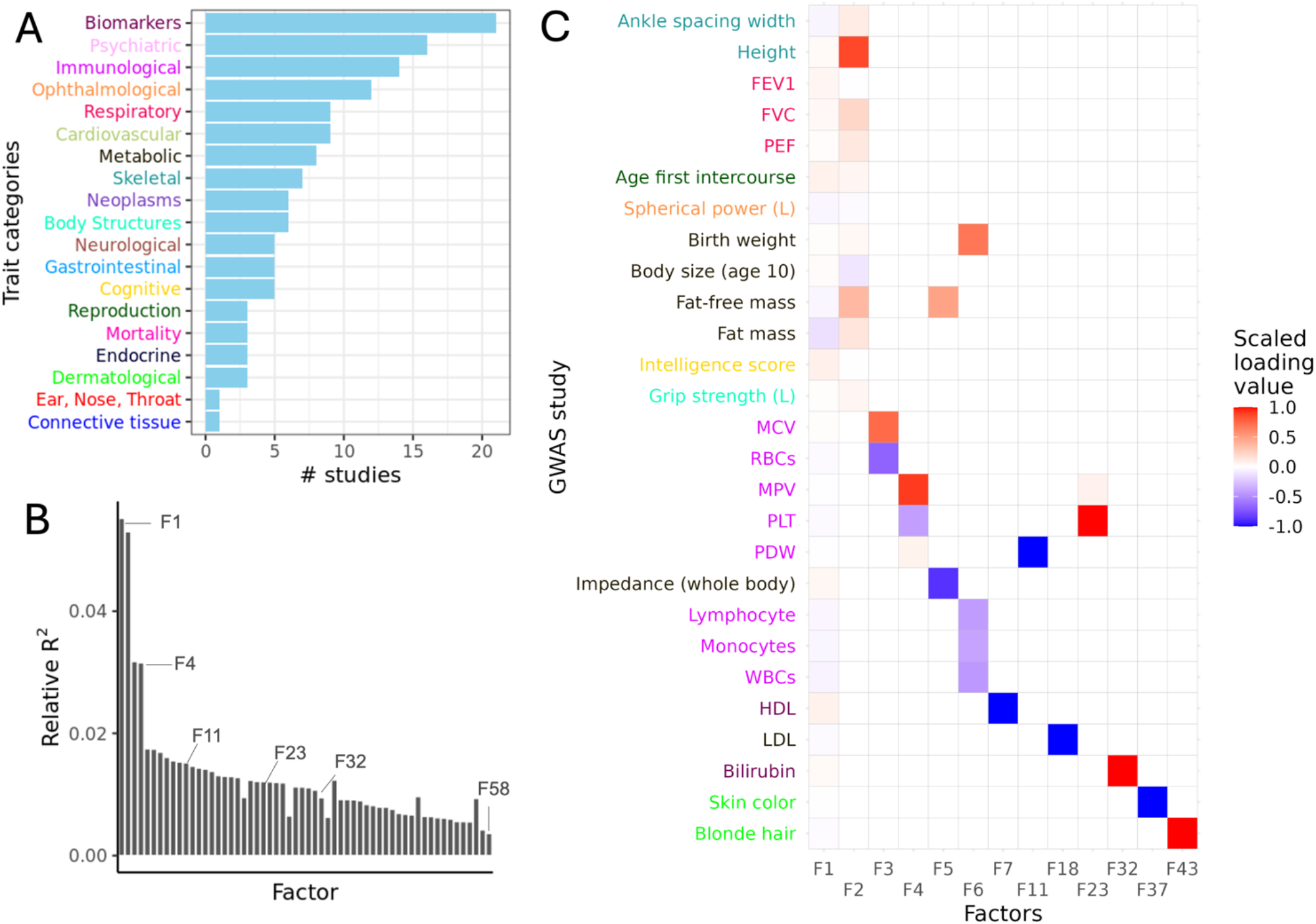
GLEANR analysis across 137 diverse European UKBB GWAS phenotypes yields 58 factors. A) Number of GWAS by phenotype category included in this analysis. Traits span 19 phenotype categories from across the human phenome. Each category label has a distinct color corresponding to member phenotype colors in (C). B) Scree plot of variance explained by 58 factors estimated by GLEANR, from F1 to F58. Factors are ordered in approximately decreasing order of data variance explained. C) Heatmap representation of selected factors in ***V*** and the traits with non-zero entries in those factors. Trait colors match the corresponding categories in panel A, and columns are scaled to have unit norm. Selected factors include the first seven explaining the largest proportion of data variance in addition to six other factors discussed in the text.

From the 137 input phenotypes, GLEANR identified 58 factors, 27 of which were loaded (e.g. had non-zero values) on only a single trait (singleton factors) and one (the uqibuitous factor, F1) was loaded on all 137 traits (Table S3). The remaining factors were loaded on up to 13 traits (mean 3.6). Each trait was weighted on between one and ten factors (median two). On average, 12.7% of SNPs per loading had values of exactly zero. Collectively, these 58 factors explained 72.6% of variance in the covariance-adjusted summary statistics (Fig 3B; a selection of 13 factors are visualized as a heatmap in 3C).

Factor 1, which explained the largest proportion of variance from input summary statistics (5.5%), was loaded on all traits (ubiquitous factor). SNP loading scores (U_1_) correlated strongly (*ρ* = −0.75) with the mean effect size per SNP across input GWAS, while trait scores (V_1_) correlated with the average genetic correlation of each input GWAS relative to all others (*ρ* = −0.80; Fig. S5B). Of all traits we analyzed, 42% loaded only on the ubiquitous factor (values of exactly zero on all other factors). These traits had fewer nominally significant SNPs included in our analysis than traits loaded on multiple factors (median of 29 SNPs per study, 1,013 SNPs per study, respectively), and were predominantly categorical phenotypes (88%). Therefore, GLEANR loads traits with few high-confidence genetic associations on the ubiquitous factor, which reflects average SNP effects across all input studies.

To test for tissue-specific regulatory effects captured by ubiquitous factor variants, we estimated heritability enrichments among chromatin tissue markers with stratified-LDSR^32,33^ (S-LDSR). Factor 1 SNPs had heritability enriched in central-nervous system related tissues (*q* = 1.74 × 10^−21^), particularly in brain markers in the fetal brain, prefrontal cortex, and germinal matrix tissues (all *q* < 1.0 × 10^−7^; Fig. S5A). Our analysis included several brain-related phenotypes (16 psychiatric, 5 neurological, and 5 cognitive traits), ten of which were loaded only on F1. This enrichment may also reflect the influence of polygenic phenotypes correlated with BMI (including whole body fat mass and whole body impedance), which has reported connections to brain gene expression markers^33–35^ and proteins^36.^ Consistent with this, gene set enrichment of top F1 genes (see Methods) in DisGeNET^37^ gene sets nominated BMI (*q* = 2.1 × 10^−8^), intelligence (*q* = 2.88 × 10^−7^), and cognition (*p* = 1.45 × 10^−4^) as top traits associated with F1 genes. Thus, the ubiquitous factor implicates brain-specific tissue markers contributing to a pathway shared broadly across the studies we evaluated.

### GLEANR factors span diverse polygenic architectures and signatures of selection

Genetic architectures vary widely across traits^31,41,42^ and regions of the genome^42^. Studies modeling GWAS effects as mixtures of multiple distributions^43,44^ provide evidence that complex traits have contributing components with different genetic architectures. We hypothesized that factors decomposing GWAS may have underlying polygenic architectures and signatures of negative selection that could meaningfully differ from their composite traits.

To characterize these properties, we first estimated factor polygenicity using a previously reported measure of the effective number of associated SNPs^41^. Scores are reported as a proportion of all SNPs under evaluation, with larger scores indicating a larger number of SNPs contributing to the factor^41^ (Fig. 4A, x-axis). Reported estimates reflect our ascertainment of pleiotropic variants in heritable traits, so are not directly comparable to genome-wide scores. F1 is among the most polygenic factors, with a broad effect size distribution and fewer than 8% of SNPs with a weight of zero. Other top polygenic factors (F52, F57, and F58) are loaded on neurological and behavioral traits including difficulty sleeping, narcolepsy, fluid intelligence score, smoking, and chronic pain. This is consistent with reported findings that behavioral and cognitive phenotypes are highly polygenic^31,41^. Factors with lower estimated polygenicity include singleton factors loaded on low-density lipoprotein (LDL; F18), and bilirubin (F32).

**Figure 4:**
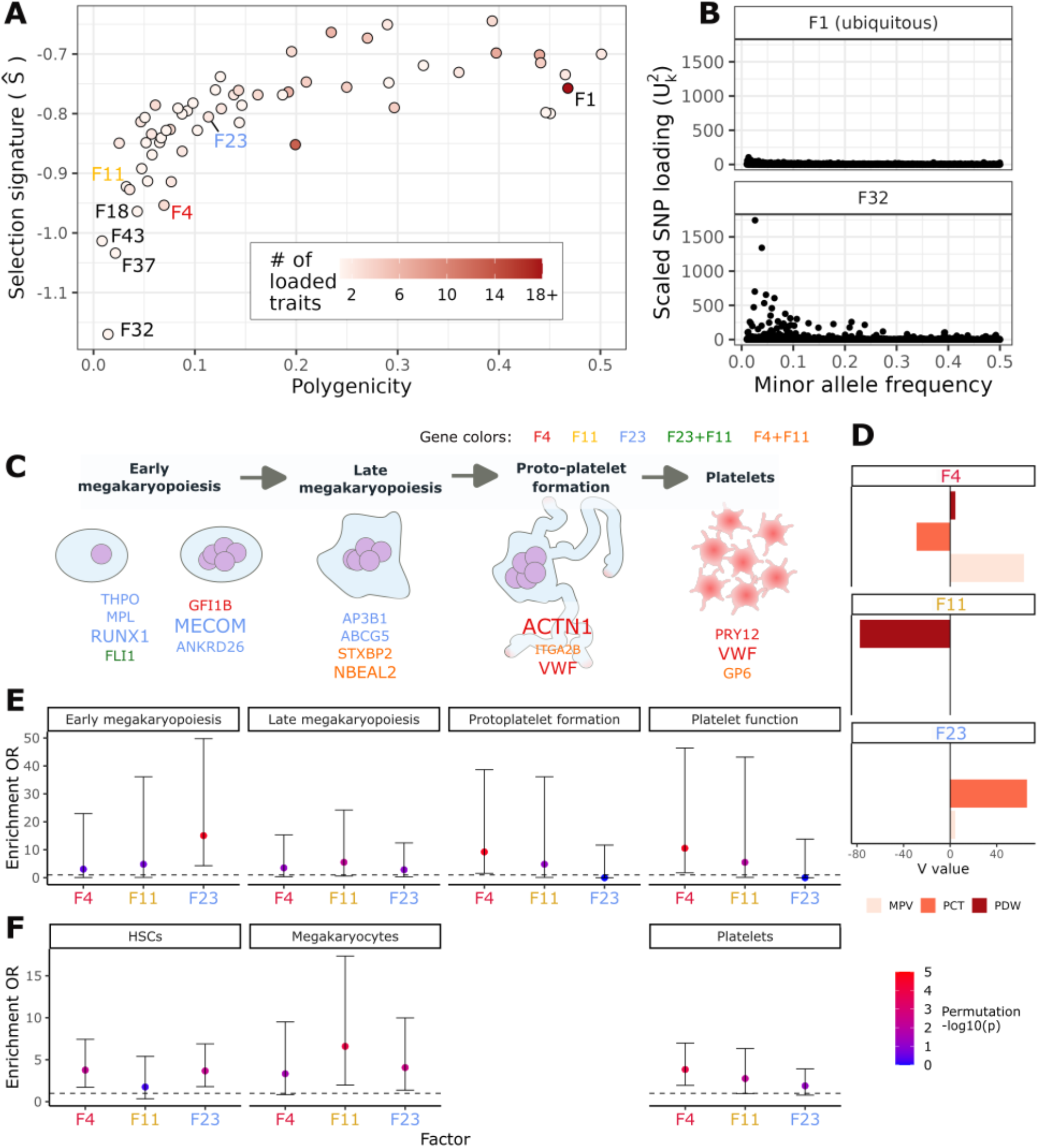
GLEANR factors reflect diverse genetic architectures and distinguish gene activity at specific stages of platelet formation. A) Relative polygenicity (x-axis) and MAF-effect size relationship (*S*_*k*_, y-axis) per factor. Points are colored by the number of traits loaded on each factor and labelled factors correspond to those referenced in the text. B) Relationship between SNP factor effect size and minor allele frequency on F1 (weak selection signature) and F32 (strong selection signature). ***U*** effects have been scaled to have unit variance for visualization. C) Schematic representing stages of megakaryopoiesis and platelet formation, adapted from Lentaigne *et al*.^38^ and Collins *et al*.^39^ Top factor genes associated with inherited platelet disorders are listed. Gene color indicates the factor in which it is a top factor gene and size indicates the average frequency with which it is nominated as a top factor gene. D) Trait weights for mean platelet volume (MPV), platelet count (PCT), and platelet distribution width (PDW) on factors 4,11, and 23. E) Enrichment of top factor genes in inherited platelet disorder genes at stages of platelet formation. Bars represent 90% confidence intervals and color indicates significance in permutation enrichment test. F) Cell type enrichment from PanglaoDB^40^ cell-type gene signatures for each factor in hematopoietic stem cells (HSCs), megakaryocytes, and platelets. These cell types were the top enriched for F23, F11, and F4, respectively.

Second, we quantified the relationship between factor effect sizes and SNP allele frequencies to examine signatures of negative selection^31,45–47^. In this model, the variance of factor SNP weights are a function of MAF (previously reported as the *α*-model^46^), which we report with the parameter *S*_*k*_^31^. Large negative values of *S*_*k*_ correspond to large effect variants enriched at low allele frequencies, often interpreted as a signature of negative selection^31,46^. F1 had the second-smallest *S*_*k*_ among all factors (*S*_6_ = −0.76), while F32, which had minimal polygenicity, reflected the strongest selection signature (*S*_:5_ = −1.17; Fig 4B). While all *S*_*k*_ estimates were negative, we emphasize that large *S*_*k*_ magnitudes likely reflect the ascertainment of the included variants, which all have nominal genome-wide significance in at least one phenotype. Thus, we limit the interpretation of *S*_*k*_ to a relative scoring among factors.

The four factors with the largest *S*_*k*_ magnitudes were all singleton factors, loaded on total bilirubin (F32), skin color (F37), blonde hair (F43), or low-density lipoprotein (LDL, F18). Factors 32 and 18 are most prominently enriched for heritability in liver-specific chromatin markers (S-LDSR tissue-adjusted factor q = 0.00026 and 0.00092 respectively; Table S4), consistent with the role of liver in cholesterol metabolism. We tested top factor genes corresponding to lead factor variants (matched using the Open Targets platform^48,49^, Table S5) for KEGG pathway enrichments^50^ (see Methods). F18 genes were enriched in cholesterol metabolism, fat digestion and absorption, and glycerolipid metabolism, reflecting cholesterol production and recycling pathways (Table S6). Top F32 genes were enriched in bile secretion and glycerolipid metabolism pathways, with top SNPs near UGT genes involved in bilirubin metabolism^51^, capturing biology related to cholesterol elimination through the bile (Table S7). F18 and F32 were minimally correlated (*ρ* = 0.04), corresponding to distinct components of lipid metabolism with evidence of negative selection, consistent with previous reports of a strong selection signature for dislipidemia^31^ in genomes of European ancestry.

We also observed large *S*_*k*_ estimates on F37 and F43 (singletons for skin color and blonde hair, respectively), consistent with prior positive selection scans on European ancestry genomes ^52,53^. Pigmentation genes under selection^52,53^, including *OCA2, HERC2, SLC45A2, TYR*, and *TYRP1*, were among top factor genes in both of these factors. Overall, GLEANR identifies genetic factors with prominent allele frequency-effect size relationships that reflect genetic loci previously reported in selection scans.

Factors with *S*_*k*_ of greater magnitude were typically loaded on few traits, suggesting that latent traits under strong negative selection may be less likely to share genetic effects with other phenotypes. These stronger MAF-effect size signatures typically appeared in less polygenic architectures (Fig. 4A and 4B). While the observed inverse relationship between factor polygenicity and *S*_*k*_ is consistent with published complex trait architecture metrics,^31^ at extremes of the polygenicity spectrum (Fig. S6), we do not detect factors with both low polygenicity and weak signatures of negative selection. This is likely due in part to our filters for selection of heritable phenotypes and pleiotropic SNPs as input to GLEANR, potentially precluding detection of factors with few causal variants bearing weak effects. Overall, GLEANR decomposes traits into factors spanning a range of polygenic architectures and signatures of selection.

### Factors weighted on platelet traits distinguish stages of platelet formation

GLEANR factors across UKBB GWAS identified instances of disease-relevant biology shared across traits. Our analysis included three platelet-related phenotypes (mean platelet volume (MPV), platelet count (PCT), and platelet distribution width (PDW)) which were decomposed into three distinct factors (F4, F11, F23; see Fig. 4D), in addition to the ubiquitous factor (F1). To characterize differences between F4, F11, and F23 (henceforth, “platelet factors”), we closely examined their SNP loadings, genetic architecture profiles, and top factor gene enrichments. As expected, top variants nominated by these factors corresponded to GWAS associations for these phenotypes^48^, with strong evidence for regulatory roles as expression quantitative trait loci (eQTLs) in whole blood (Table S8). For instance, rs2228367, the second lead variant in F4, is a replicated GWAS hit for all three phenotypes^24,54,55^ and an eQTL in whole blood in multiple studies^56,57^ for tropomyosin 4, an actin-binding protein with well-documented effects in platelet diseases.^58,59^

Among platelet factors, F4 had the strongest selection coefficient estimate (Ŝ = 0.95; fifth largest magnitude in the entire analysis). Consistent with negative selection acting on large effect SNPs, top F4 SNPs had larger average effect size magnitudes across MPV, PCT and PDW than F11 (Wilcoxon rank sum test, *p* = 5.5 × 10^−6^) and F23 (*p* = 4.6 × 10^−79^) SNPs (Fig. S7). Given this selection signature, we speculated that variants captured by F4 might adversely influence human fitness.

We also observed that PCT and MPV had opposite directions of effect in F4 (Fig. 4D), suggesting that F4 SNPs drive platelet volume and platelet count in opposing directions. This is consistent with previously reported negative correlation between GWAS effects for these traits^39^ and with negative correlation of the traits themselves across patients^60,61^. Published work has hypothesized that alleles contributing to both lower PCT and elevated MPV could explain hereditary cases of the blood condition macrothrombocytopenia^39^. Hereditary macrothrombocytopenia (HMTP) is characterized by elevated platelet size (>12 femtoliters) and low platelet counts (< 150,000/µL)^62^ and is associated with many other blood-related diseases (e.g. Sitosterolemia [MIM:210250], Grey Platelet Syndrome [MIM:139090], Bernard-Soulier syndrome [MIM: 231200], platelet-type bleeding disorders [MIM:187900,616937], and cancer susceptibility [MIM:616216])^39^. While previously thought to be a Mendelian condition, growing evidence from GWAS suggests polygenic contributions to both HMTP and other blood diseases ^39,55^.

To evaluate whether F4 captured genetic effects contributing to HMTP, we tested top F4 genes for enrichment in known HMTP-causing genes ^39^ (Table S9). Given that multiple SNPs were mapped to the same gene in our analysis, we considered only unique genes in our enrichment tests and used random subsamples of our background gene set to generate an empirical null reflecting this imbalance (Methods). We found that F4 was strongly enriched for HMTP genes (OR = 6.5, empirical p = 1e-04), as were F11 and F23 (OR 6.1 and 6.3, respectively), suggesting a possible source of differential fitness that may impact the selection signature of F4.

Next, we sought to explore the cell-type and context specificity of the platelet factors, noting that disease-relevant processes can occur in early and transient stages of cellular differentiation^63–65^ in addition to terminal and adult cell types^56,66,67^. We tested each factor for enrichment in genes known to cause inherited platelet disorders at distinct stages of platelet formation^38,39^, specifically during early megakaryopoiesis, late megakaryopoiesis, proplatelet formation, and mature platelet function (Fig. 4C, Table S9). We observed that F23 was strongly enriched for inherited platelet disorder genes in early megakaryopoiesis (corrected empirical q<0.00008) and was the only factor to capture thrombopoietin (*THPO*), the critical growth factor promoting megakaryopoiesis^68^, as a top gene. F11 related to late megakaryopoiesis (q = 0.02), and F4 reflected the later stages of proplatelet formation (q< 0.00008) and mature platelet function (q<0.00008; Fig. 4E).

The correspondence between factors and different stages of platelet formation was further supported by complementary cell-type and GO-term enrichments. In an enrichment test of top factor genes in cell type gene markers from the PanglaoDB^40,69^, the strongest detected enrichment for F23 was in hematopoietic stem cells, while F11 was most strongly enriched in megakaryocyte gene markers, and F4 in platelets (Fig. 4F), though these enrichments are no longer significant after adjusting for multiple hypothesis testing across all cell types (Table S10).

Consistent with extensive transcriptional regulation during early megakaryopoiesis, F23 had top GO molecular function enrichments in DNA binding (BH-corrected q=0.005) and transcription factor binding (q=0.04) and had GO biological process enrichments in myeloid cell development (q=0.03), myeloid cell differentiation (q=0.08), and hemopoiesis (q=0.03) (Table S11). Capturing the cytoskeletal dynamics involved in proplatelet formation^39,70^, F4, had top GO molecular function enrichments including tubulin binding (q=0.03), cadherin binding (q=0.03), and microtubule binding (q=0.05, Fig. S8, Table S12). Our observations across these platelet factors provide strong evidence that sparse factorization of GWAS by GLEANR can decompose trait effects into interpretable components that capture distinct disease-relevant cellular processes.

## Methods

### GLEANR Model

GLEANR operates on marginal GWAS summary statistics, with effect size estimates in the matrix ***B*** ∈ℝ^*N*×*M*^ and standard error estimates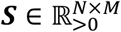. SNPs are indexed by *n* ∈ [1, … *N*] and traits by *m* ∈ [1, … *M*]. Typically, substantial covariance is observed across both columns and rows due to sample sharing among traits and linkage disequilibrium between SNPs, respectively. To account for the impact of linkage disequilibrium, GLEANR requires SNPs under evaluation to be approximately uncorrelated, selected by an appropriate pruning or clumping procedure (e.g. r^2^ < 0.3 across 250KB windows). We model the covariance associated with GWAS estimation error and sample sharing as follows:

Let *β*_***n***_ be a single row of ***B*** and ***s***_***n***_ a single row of ***S*** along SNP *n*, where:

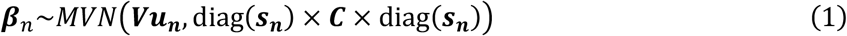

***V*** ∈ℝ^*M*×*K*^ maps *M* related traits onto *K* latent factors, so traits sharing a pleiotropic genetic factor have non-zero values along the same column of ***V. u***_***n***_ ∈ℝ^*K*×1^ (a single row of the matrix ***U*** ∈ℝ^*N*×*K*^) is a sparse vector indicating the contribution of variant *N* to each of K latent factors. We allow ***U*** and ***V*** to be signed, as the parity with which a SNP contributes to a phenotype association (i.e. if it is a risk or protective SNP) is dictated by the arbitrary measure of traits. diag(***s***_***n***_) ∈ℝ^*M*×*M*^ contains standard errors for SNP *n* across all studies along the diagonal, and ***C*** ∈ℝ^*M*×*M*^ is a block correlation matrix of non-genetic confounding covariance structure between GWAS.

GLEANR imposes sparsity constraints on both ***U*** and ***V*** such that 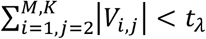 and 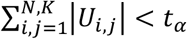, where *t* is a threshold corresponding to the lasso regularization weights for ***V*** and ***U*** (*λ* and *α* respectively, see below).

#### Model fitting procedure

GLEANR model fitting begins by initializing ***V*** to the first *K*_*init*_ − 1 right singular vectors of ***B***. An additional column is appended to reflect the dominant direction of correlation across all studies, with signs corresponding to the first eigenvector of cor(***B***). We perform alternating iterations of generalized least squares regression with L1 regularization to update 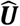 by regressing 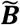 against 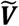, where the tilde indicates elements of the matrix have been transformed to adjust for GWAS estimate uncertainty and confounding correlation (rows of 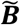 are 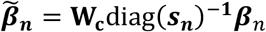 and 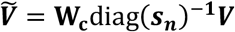 We then regress 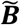 against 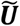 to update 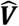 (where elements of 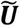 are 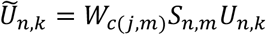 expanded over all *j, m* ∈ [1, … *M*]; Supplemental Note 1). We repeat alternating regressions until our objective function achieves convergence (Equation 2):

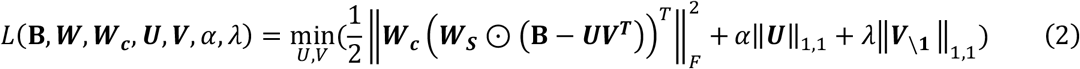

***W***_***S***_ is a matrix of weights with elements 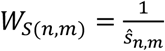 and ***W***_***c***_ ∈ℝ^*N*×*M*^ is a decorrelating transformation derived from 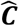 (see below). Here, 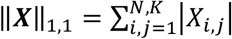, is the entry-wise L1 matrix norm. The first column of ***V*** is unregularized to retain the dominant correlation structure across traits.

This objective monotonically decreases at each iteration because it is bi-convex (Fig. S9) and achieves convergence when the change between iterations is smaller than a specified small threshold *ε*, with a default of < 0.1% of the previous iteration’s value. This procedure is implemented using sparse matrix functions in *glmnet in R 4*.*2*.3^18^ (Supplemental Note 1).

This model fitting procedure is summarized below:

##### Algorithm 1

GLEANR model fitting

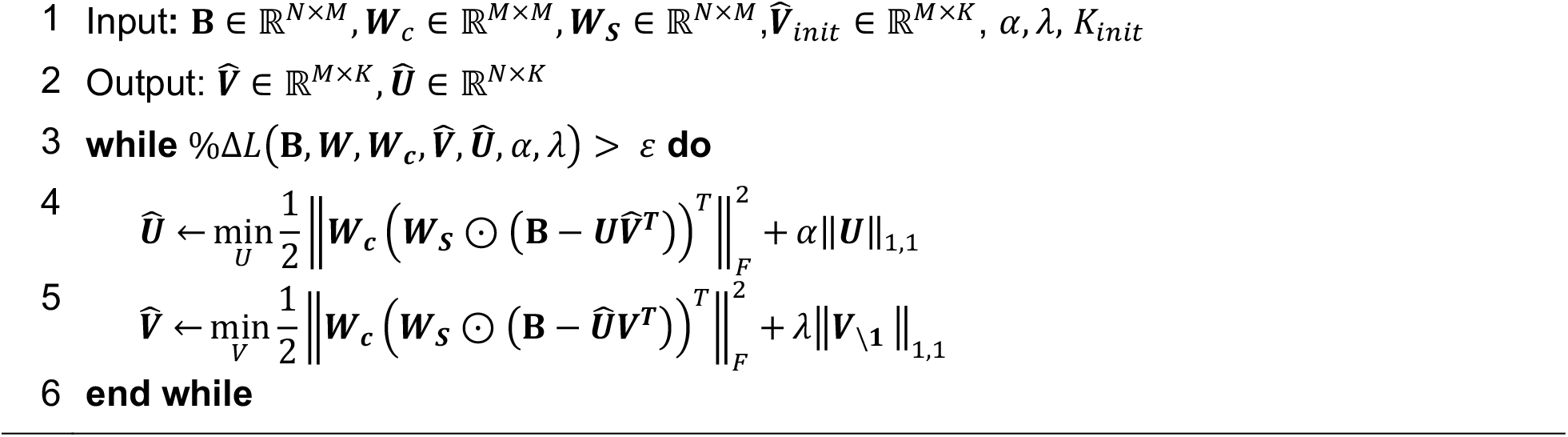

*Estimation of* ***C*** *and decorrelating transformation (****W***_***c***_)

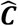 may be estimated from the intercept term of XT-LDSC^3,4^, from phenotype correlations and the number of overlapping samples^25^ (when available), or from the empirical correlation of truncated z-statistics^71,72^. In this work we favor estimates from XT-LDSC.

Empirically, GLEANR is sensitive to large elements 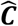 To mitigate these effects and account for uncertainty in estimates of 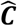, we apply shrinkage to its elements in the following steps:

1. Require 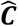 to have a block structure, reflecting our expectation that ***C*** is sparse and contains clusters of correlated traits^25^. Block elements are identified through hierarchical clustering on the matrix of cohort overlap estimates, followed by pruning branches with a cutoff corresponding to pairwise correlations ≥ 0.2.
2. Apply Wolf-Ledoit shrinkage^73^ to the block correlation matrix, adjusting for uncertainty in correlation estimates^74^ by calculating a matrix 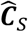:

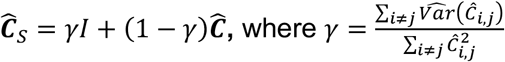

This procedure has the added benefit of yielding a theoretically positive definite matrix 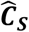 as the linear combination of positive definite and semi-definite matrices^73^.

When 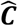 is estimated using XT-LDSR,^3^ GLEANR provides the option to scale *g*_*cov*–0*nt*_ statistics by the product of the standard deviation of input traits, to adjust for the selection of heritable traits in analysis.

### Model selection procedure

GLEANR has three model hyper-parameters: *α, λ* (which correspond to the degree of sparsity in ***U*** and ***V***, respectively), and the number of factors *K*_*init*_ at which to initialize GLEANR. Although multiple approaches for model selection exist in the context of matrix factorization^20,75–77^, we employ the Bayesian Information Criteria (BIC)^20,22,75,78^ due to its interpretability and computational efficiency. In high-dimensional problems, the BIC favors overly complex models^79,80^, so we include an additional term^79^ to adjust for models with many covariates receiving a higher likelihood under the standard prior. We calculate the BIC as:

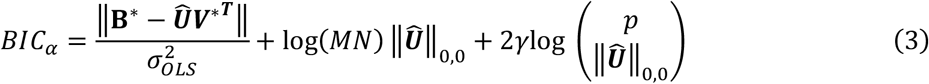

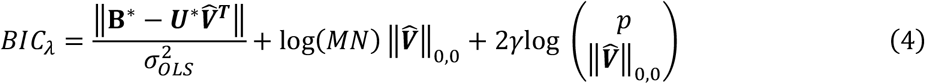

where 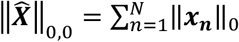 (e.*g*. the number of non-zero elements in the matrix), is an unbiased estimate for the degrees of freedom of the lasso^21^, and *γ* = 1 − lo*g* (*NM*)/2 lo*g*(*P*) is a scaling term that controls the contribution of the added BIC extension term^79^. *p* is the total number of covariates under consideration in each model comparison (*NK* for ***U*** or *MK* for ***V***). This “extended” BIC yields more consistent results across GWAS cohorts than the traditional BIC in our tests (Fig. S10).

#### *Selection of α, λ, and K*_*init*_

During GLEANR’s initial iterations, we select either the *λ* (when fitting ***V***) or *α* (when fitting ***U***) which minimizes the BIC of the current step until *L*_***B****IC*_ = *BIC*_*α*_ + ***B****IC*_*λ*_ converges, at which point *α, λ* are fixed. In a single iteration of model fitting, we fit ***U*** at 100 settings of *α* along the log-scale of possible values (from *α*_*max*_, where all of 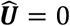, to *α*_*max*_ × 10^−4^). GLEANR selects 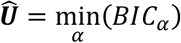, then repeats this procedure with respect to ***V*** and *λ*. Alternating iterations continue until *L*_***B****IC*_ is no longer declining (or for at least 5 iterations), at which point *α* and *λ* are fixed at 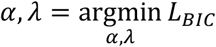 and GLEANR proceeds with model fitting (see Algorithm 1).

In practice, we observe that *L*_***B****IC*_ typically converges in fewer than 10 iterations (Fig. S11).

To select *K*_*init*_, GLEANR performs an adaptive grid search to select the *K*_*init*_ which minimizes *L*_***B****IC*_ at convergence (Supplemental Note 1). GLEANR first samples *K*_*init*_ from quantiles of the range [1, *M* − 1], then tests additional settings in the neighborhood of 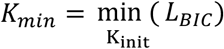, halfway between previously tested values. This procedure continues for a specified number of search points (default =10) or until *K*_*m*i*n*_ no longer changes.

##### Algorithm 2

GLEANR model selection

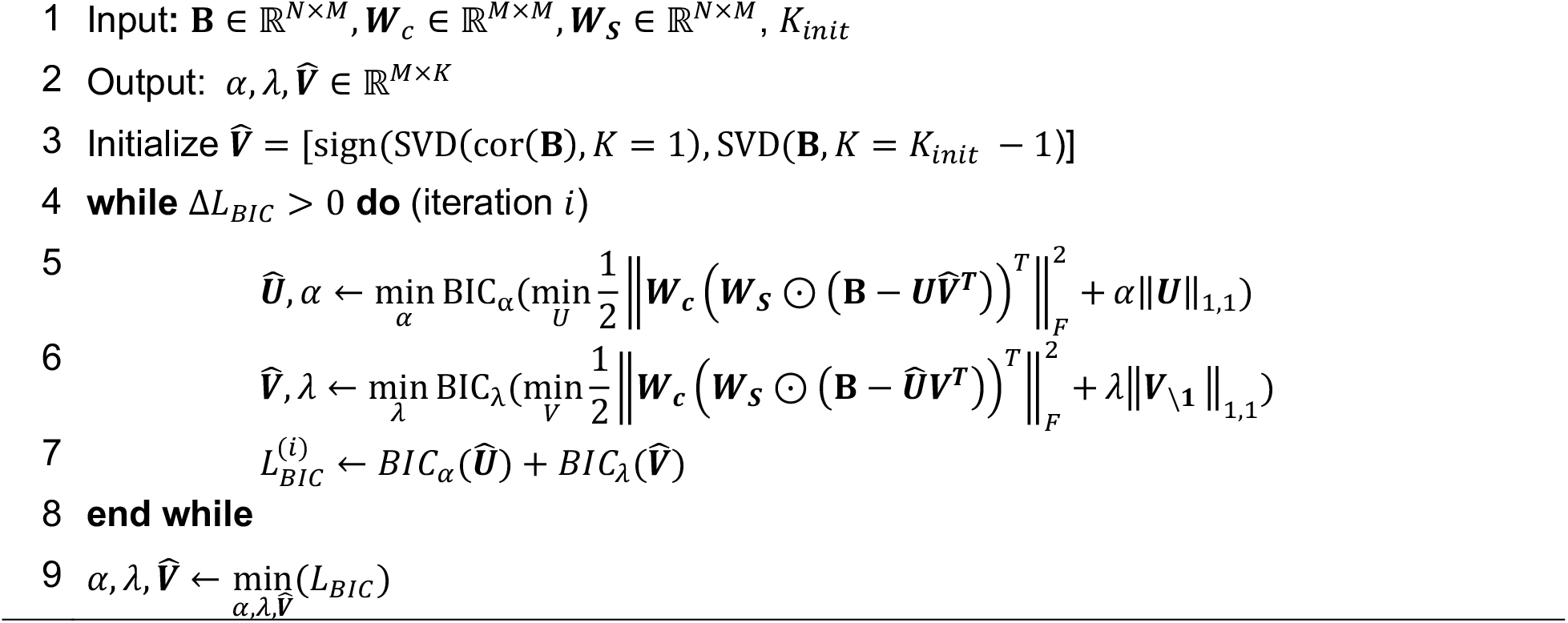

### Simulations

Our study included two simulation frameworks: 1) to isolate the impact of sample sharing on the detection of spurious genetic factors and 2) to test reconstruction of known genetic factors. In (1), we directly specified the degree of sample sharing when estimating summary statistics. In (2), we simulated a latent basis (***U, V***) and added noise corresponding to covariance structure induced by sample sharing between studies.

#### 1) Direct simulation of GWAS summary statistics

Three populations of 91,000 individuals were simulated, with genotypes for 1000 SNPs sampled from Bernoulli distributions for maternal and paternal alleles. Count probabilities came from minor allele frequencies of randomly selected pruned SNPs nominated by UKBB studies in our comparative UKBB-FinnGen analysis. Phenotypes were modeled based on the height, weight, and BMI of 10,000 UKBB individuals. Population parameters (height mean and variance and weight predicted from height) were used to generate 3 random phenotypes as; *p*_1_ ∼ *N*(169.9, 86.5), *p*_2_ = 0.93*p*_1_ − 80 + *ϵ*_5_, *ϵ*_5_ ∼ *N*(0,182.3) and 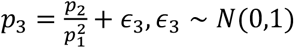 We selected 30,000 individuals from each cohort per phenotype and estimated GWAS summary statistics with linear regression on standardized phenotypes, such that no individual was used more than once (for a total of nine sets of summary statistics). We repeated this procedure, reusing samples within a cohort at levels of 10,15,20,23,26, and 30-thousand repeated samples. We considered all unique pairwise combinations of overlap levels across the three cohorts for a total of 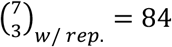 overlap settings, repeated 10 times each (Table S13). The percent of overlapping samples we report in Figure 2 reflects total number of samples which were used multiple times when estimating GWAS from the three cohorts, divided by the maximum number of possible shared samples (90,000).

We performed matrix factorization and model selection on each of these 840 sets of summary statistics using the singular value decomposition (SVD), flash, and GLEANR, as described in Table 1. SVD was implemented using the BiocSingular *runPCA* function. “flash,”^26,81^ a robust and flexible method for variational Bayesian matrix factorization, was implemented using the “flashier”^81^ package with backfitting to update factors after initial fitting and all other settings as default. GLEANR without covariance adjustment is equivalent to standard GLEANR with ***W***_***c***_ = ***IM***,the *M* × *M* identity matrix. To estimate 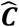, we used the known simulation sample overlap and empirical phenotypic correlation, with off-diagonal elements given by 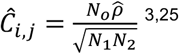 and standard error estimates 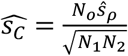, with 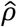 and 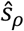 from the R *stats*: : *cor. test* utility and *N*_*o*_ indicating the number of samples shared between two studies. Performance was assessed by counting the number of non-zero factors nominated by each model selection procedure.

**Table 1:** Summary of methods evaluated in direct GWAS-based simulations and their corresponding inputs. Matrix names (***B, S, W***_***S***_, ***W***_***c***_) match those in the main text.

| MF method | Model selection method (to select K) | Input |
| --- | --- | --- |
| SVD | Factors above the “elbow” of the scree plot | Z-scores ($Z = B \odot W_S$) |
| SVD | Factors eigenvalues greater than the average eigenvalue: $\{k_\lambda: \lambda_k > \frac{1}{M} \sum_{k=1}^M \lambda_k\}$ | Z-scores ($Z = B \odot W_S$) |
| SVD-adj | Factors above the “elbow” of the scree plot | “Decorrelated” Z-scores<br>$Z_d = (W_c(B \odot W_S)^T)^T$ |
| SVD-adj | Factors eigenvalues greater than the average eigenvalue: $\{k_\lambda: \lambda_k > \frac{1}{M} \sum_{k=1}^M \lambda_k\}$ | “Decorrelated” Z-scores<br>$Z_d = (W_c(B \odot W_S)^T)^T$ |
| flash | “Greedy” addition of non-zero factors, followed by “backfitting” procedure | $B, S$ |
| GLEANR (no adjustment) | Grid search for $K_{init}$ followed by regularized model fitting | $B, W_S$ |
| GLEANR | Grid search for $K_{init}$ followed by regularized model fitting | $B, W_S, W_c$ |

#### 2) Simulation of latent genetic basis

To evaluate latent space estimation in the presence of confounding covariance, we simulated summary statistics from genetic factors and added noise reflecting sample sharing across studies. We simulated effect sizes (***B***) for 100 SNPs across 10 traits as ***B*** = ***UV***^***T***^***a***, with prespecified ***U, V*** and ***a*** a vector to scale causal effect sizes to match variance of previously reported continuous trait phenotypes^43^. We generated three each of ***U*** and ***V*** with *K* = 5 for a total of nine distinct ***B*** matrices. Elements of ***U*** were sampled (at *k* factors, *n* SNPs) from

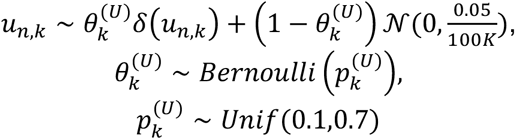

Elements of ***V*** (at *k* factors, *m* traits) were drawn from:

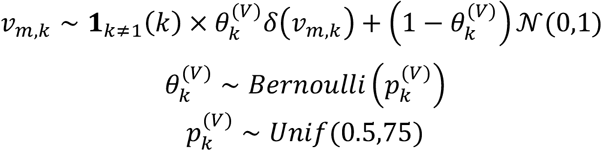

where **1**_*k* ≠ 1_(*k*) is an indicator function returning zero for *K* = 1 (the first factor), and *δ*(·) is a point mass at zero. All columns of ***U*** and ***V*** were required to have at least one non-zero element. We then added noise ***E***, where a single row is given by ***ϵ***_0_ ∼ *MVN*_*M*_(0, diag(***s***_***n***_) × ***C*** × diag(***s***_***n***_)) Elements of the vector ***s*** are given by 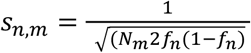, where *f*_*n*_ (minor allele frequency of SNP *n*) was sampled from the European UKBB variants used in our analysis of 137 diverse phenotypes. *N*_*m*_ (sample size of study *m*) was simulated at levels of 5,10,50,100 and 200 thousand samples for each study, and one scheme with a “mixed” number of samples (10-200 thousand).

***C*** was simulated with elements

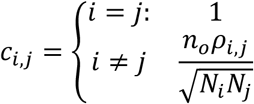

where *N*_*i*_ corresponds to the total sample size of study *i. n*_*o*_ and *ρ*_*i,j*_ each correspond to one of three different cohort overlap structures tested. In the case of no overlap (Structure 1), *ρ*_*i,j*_ = *n*_*o*_ = 0. For block cohort overlap structures (Structures 2-3), we let *n*_*o*_ = 0.9 × min (*N*_*i*_, *N*_j_) and *ρ*_*i,j*_ = 1 for all diagonal entries. In the “single block” (Structure 2) setting, *ρ*_*i,j*_ ∼ *Unif*(0.75,0.99) for traits 4-8, and zero for all others. In the “double block” setting (Structure 3), *ρ*_*i,j*_ ∼ *Unif*(0.4,0.7) for traits 1-5 or 6-10, zero otherwise. These resulted in a total of 162 different evaluation settings, each of which had ***E*** resampled ten times.

Each method was specified to yield K=5 (or a maximum of *K* = 5 in the case of GLEANR and flash). In addition to the methods above, we included flash with the Kronecker variance setting, which models a block matrix covariance structure, and the GWAS-specific variational method FactorGo^8^.

### Matching factors and evaluating factorization performance

To determine how closely the 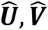 predicted by factorization methods matched simulated matrices, we evaluate squared correlation (R^2^) across all predicted factors. Matrix factorization solutions are not guaranteed to be identifiable, and different solutions may reflect equivalent matrix structure that differ by an arbitrary matrix rotation and scaling. To address this, we found the factor order (***I***_***O***_) and sign vector (***μ***) that maximized the global correlation across both matrices by optimizing the following:

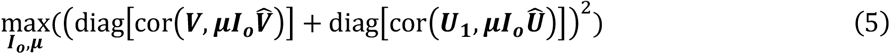

where **I**_**O**_ is an orthogonal rotation matrix with exactly one non-zero element per column of value 1, ***μ*** is a vector of signs with elements *μ*_*i*_ ∈ [1, −1], and cor(**X**_**1**_, **X**_**2**_) is the *K* × *K* correlation matrix between columns of **X**_**1**_ ∈ℝ^*N*×*K*^ and **X**_**2**_ ∈ℝ^*N*×*K*^. When the dimension of predicted matrices was fewer than the true matrices, columns were padded with zeros (Supplemental Note 2). We then calculated R^2^ across all factors, each scaled to have unit norm.

### FinnGen and UKBB cross-cohort analysis

#### Study selection

We compared GLEANR factors across 42 phenotypes from GWAS evaluated in the FinnGen^29^ and UKBB^14^ cohorts (Table S1). The FinnGen study is a large-scale collaborative genomics initiative that has analyzed over 500,000 Finnish biobank sample and has published summary statistics for over 2000 phenotype endpoints^29^. Similarly, the UKBB is a large-scale biomedical database with genetic and health information for over 500,000 UK volunteer participants aged 40-69 years^14^. Summary statistics for 41 disease phenotypes in the UKBB cohort were accessed from the PheWeb^30^ server, an online resource of GWAS in over 1400 binary phenotypes from white UKBB participants. For the remaining continuous phenotype, summary statistics came from the Neale Lab GWAS portal^24^ round 2 results (Web Resources). FinnGen and PheWeb GWAS were generated using SAIGE^82^, and all studies were controlled for sex, age or birth year, and multiple genotype PCs. To select traits for evaluation, we first manually matched phenotypes available in both cohorts. These were filtered to GWAS with at least 25 minor alleles among controls for SNPs with MAF > 0.01, an effective sample size > 5000, and one reported GWAS hit in the FinnGen GWAS manifest. Subsetting to HapMap3^28^ SNPs, we performed XT-LDSR^3^ to estimate the genetic correlation between cohorts on the 77 remaining traits, keeping only those with *r*_*g*_ > 0.8 at a Bonferroni-adjusted *p* < 0.01. From the remaining 42 studies, we selected SNPs for evaluation by filtering out ambiguous and multi-allelic variants and those in the HLA region, keeping only those with *p* < × 10^−4^ in at least one GWAS. 14750 SNPs passed these filters in both the FinnGen and UKBB GWAS, which we then pruned using plink2^83^ (–*indep-pairwise 250kb 0*.*2)* and a European 1000 Genomes^84^ LD reference panel. The remaining 1663 SNPs were input to GLEANR analysis, evaluated at *K*_*init*_ = 41. We estimated cohort-specific sample sharing from XT-LDSR,^3^ scaling reported *g*_*cov*–*int*_ statistics to adjust for the selection of heritable traits.

To assess the concordance of factors between cohorts, we calculated the Pearson correlation coefficient across all unit-scaled factors, matching factors as described above (Equation 5), with those from FinnGen GWAS used as reference factors. We tested the difference between these correlation estimates using Fisher’s Z-test for independent groups as implemented in the *cocor* R package^85^. To compare reconstructed effect sizes 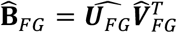 and 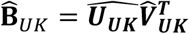 we calculated the relative root mean squared error (RRMSE):

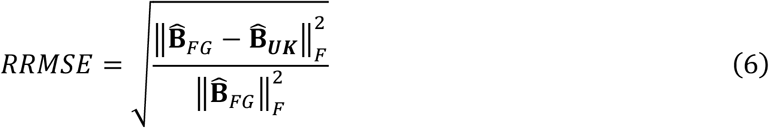

### UKBB analysis across 137 diverse GWAS

#### Study selection

Traits were selected from 7228 phenotypes with GWAS published by the Pan-UKBB^27^ project. Summary statistics were generated in a standardized pipeline across six continental ancestries (including up to 420531 European individuals; see Web Resources) using the SAIGE^82^ software. We considered for inclusion those traits passing Pan-UKBB recommended filters (valid LDSR^86^ *h*^2^ ∈ (0,1) with a significant Z-score, normal *λ*_*g*,c_ across top three powered ancestries, an LDSR inflation ratio < 0.3), studies with effective sample size > 5000, and LDSR^86^-estimated *h*^2^ > 0.05. We limited GWAS of continuous phenotypes to those performed on inverse rank normal transformed values and removed phenotypes in categories difficult to interpret or not of biological interest (e.g. activities, environment, nutrition, social interaction, parental age, and those related to hospital admission or treatment specialties). Category assignments were extended from previously published domain assignments^5^ integrated with UKBB coding information. Of the remaining traits, we added studies to our set in order of decreasing heritability, requiring that each had a pairwise *r*_*g*_ < 0.7 (estimated by XT-LDSR^3^) with all other studies in our set. This yielded 137 traits in total (Table S2, Supplemental Note 3).

#### SNP selection

From ∼29.0 million variants evaluated by the Pan-UKBB^27^, we considered biallelic SNPs designated as “high quality” (e.g. PASS variants in gnomAD and consistent frequencies across gnomAD ancestries), with INFO > 0.9, MAF > 0.01 in Europeans. From these 7.9 M variants, we selected HapMap3^28^ SNPs not in the HLA locus with GWAS *p* < 1 × 10^−5^ in at least 1 study, then LD-clumped these variants with plink,^87^ using a 1000 Genomes^84^ European genotype reference panel. During clumping, SNPs were prioritized according to the number of studies in which they achieved our nominal significant threshold, favoring the inclusion of more pleiotropic SNPs and yielding a total of 41260 SNPs (Supplemental Note 3).

#### UKBB GLEANR analysis

We performed GLEANR on the selected GWAS summary statistics using an adaptive grid search at eight settings of *K*_*init*_, selecting *K*_*init*_ = 134 which yielded the smallest *L*_***B****IC*_ score. GLEANR model fitting converged in 34 iterations, following seen iterations of model selection. 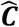 was estimated using XT-LDSC^3^ and scaled to adjust for the selection of heritable traits.

#### GLEANR factor evaluation

##### Factor tissue-marker enrichment

To test for heritability enrichment among tissue-specific chromatin markers in GLEANR factors, we performed stratified-LDSR^32^ using 489 tissue-specific chromatin markers,^33^ LD scores, and weights provided by the authors (see Web resources). GLEANR loading scores (columns of 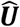) were scaled into Z-scores for input into S-LDSR, where SNP *n* of factor *k* has value

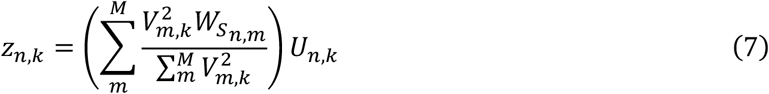

Factor sample sizes *N*_*K*_ were given as the weighted average of the sample sizes of traits in a given factor (***v***_***k***_), with 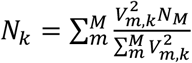. Because only 41260 SNPs were available for regression, we increased the number of LDSR jackknife blocks to 5000 (default 200) to improve standard error estimation.

To correct for multiple hypothesis testing, we performed Benjamini-Hochberg (BH) adjustment across all enrichment tests per factor and reported BH-adjusted p-values as “q-values”^88^ (Table S4). To evaluate which tissue category yielded the most prominent enrichment across all 489 tested chromatin markers per factor, we further adjusted for the unequal number of markers per tissue category. To do this, we performed Bonferroni adjustment of p-values for all factor tests within a given tissue category, retained only the enrichment with the smallest p-value per group, and then performed BH adjustment across the lead tissue-adjusted p-value from each category.

##### Factor genetic architecture

We estimated the polygenicity and the selection signature parameter^31,46^ for each factor. To estimate the polygenicity of factor *k*, we leveraged the kurtosis (scaled 4th moment) of SNP effect sizes^41^, reporting 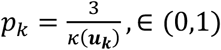, where ***k***(***u***_***k***_) is the empirical kurtosis of elements of ***u***_***k***_. Values close to one reflect larger tails^89^ on the effect size distribution and indicate higher polygenicity.

To estimate selection coefficient *S*_*k*_, we modeled non-zero factor effect sizes as 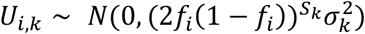, where *f*_*i*_ is the MAF at SNP *i*. Using R’s *optim* function, we find the *S*_*k*_ and 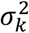 maximizing the empirical log-likelihood of non-zero effects in factor ***u***_***k***_.

##### Top SNP selection

We defined “top factor SNPs” as those in the top tertile above the “elbow” of ranked factor SNP effect magnitudes. For factor loading ***u***_***k***_, we ranked SNPs according to their magnitude in decreasing order, and then selected the SNP with effect size magnitude (i.e. |*U*_*n,k*_|) at the graphical “elbow” of this distribution (i.e. achieving maximal distance from a line between the largest and smallest magnitude SNP effect size). We designated SNPs in the top third of effect size magnitudes above this elbow as top factor SNPs. The number of SNPs per factor chosen in this manner correlated strongly with estimates of polygenicity (*ρ* = 0.85; Fig. S12).

##### SNP-gene mapping and gene set enrichment analysis

All SNPs were mapped to a putative candidate gene using the latest variant-to-gene mapping from the Open Targets^48,49^ platform (download version 2022-10-11). If multiple genes were assigned equal scores by Open Targets or no gene assignment was available, we selected the gene with the closest protein-coding transcript start site to the SNP.

To perform gene set enrichment on factors, we designated the set of distinct genes mapped to by top factor SNPs as the set of top factor genes. All distinct genes mapped to by a SNP in the analysis constituted our background set, yielding 10,512 unique genes (from 41259 candidate SNPs), and factors each had an average of 414.8 unique top genes.

Enrichment testing in canonical gene set libraries (including GO Biological Process, GO Cellular Component, GO Molecular Function, DisGeNET, KEGG pathways, GWAS Catalog, and Panglao DB) was performed using the *enrichr*^69,90,91^ API. Reported q-values represent p-values adjusted using the BH^88^ procedure for the number of tests per gene set library.

For gene set enrichment tests on sets related to inherited platelet disorders, genes were collected from those in previously studies ^38,39^ and grouped by the reported stage of platelet development which they disrupt in disease. Top factor genes were tested for enrichment against background genes using a Fisher’s exact test. To account for the small gene set size and the non-uniform inclusion of genes in our analysis (i.e. many genes are mapped to by multiple SNPs), we generated an empirical null distribution of enrichment statistics. For factor *k* with *N*_*k*_ top factor SNPs, we randomly subsampled *N*_*k*_ SNPs and performed a Fisher’s exact test on the distinct set of corresponding genes. This procedure was repeated 50,000 times to generate an empirical p-value where 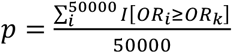 the number of tests with an odds ratio as or more extreme than the odds ratio from the true top factor genes), which we BH-corrected^88^ for the number of tests performed across factors and gene sets. A similar permutation procedure with 10,000 samplings was used to validate top PanglaoDB^40^ cell-type enrichments reported in Figure 4.

## Discussion

In this work we present GLEANR for the identification of genetic factors through sparse matrix factorization of GWAS summary statistics. GLEANR offers two key advantages over existing approaches. First, it enables hypothesis-free exploration over 100s of traits robust to the effects of shared samples among the input GWAS. Second, GLEANR encourages sparsity in the estimated factors specific to the target summary statistics, yielding latent factors that are more readily interpretable than those from dense MF approaches.

As revealed through extensive simulations, we observed that existing matrix factorization methods are susceptible to reporting spurious and confounded latent genetic factors. These spurious factors occur due to non-biological influences—particularly the effects of sample sharing among the input GWAS—which increases the covariance in GWAS effect sizes simply due to shared residual noise. These spurious factors capture non-genetic signal, complicate interpretability, and reduce reproducibility across populations and biobanks. We showed that GLEANR is more robust to identifying these spurious factors than existing methods. These findings emphasize that appropriately accounting for sample sharing, even if it only influences a subset of target studies, improves the detection of replicable components and is critical to preserve the interpretability of latent genetic factors.

Across 137 diverse GWAS in the UKBB, we estimated one ubiquitous and 57 sparse factors that reflect biological pathways underlying GWAS effects. These genetic factors are a resource for distinguishing pathways shared by multiple traits and those which are trait-specific (singleton factors). Enrichments among top factor genes and in tissue-specific chromatin markers can support hypothesis generation through exploration of previously unobserved relationships among traits, such as the enrichment we observe in brain-specific chromatin markers corresponding to Factor 1, which was loaded on all traits.

We observed that GLEANR factors decompose trait effects into components with distinct genetic architectures. Variation in factor polygenicity scores highlight that the effective number of variants contributing to factors differ. This influences factor interpretation by informing how many factor SNPs should be considered when performing enrichment tests. Components with strong MAF-effect size relationships, a signature of negative selection, are trait-specific and have lower polygenicity, consistent with estimates of polygenicity from forward evolutionary simulations that model the effects of selection^31^. Our work suggests that global evaluation of trait architecture may obscure the contributing sub-architectures of distinct genetic components.

Of the 58 factors nominated by GLEANR, we identified three factors (in addition to a ubiquitous factor) loaded on platelet phenotypes. Gene set enrichment tests provide evidence that these factors stratify variant contributions to inherited platelet disorders at distinct stages of platelet formation including hematopoiesis and megakaryopoiesis, and platelet function. Published studies of platelet phenotypes report corresponding enrichments in chromatin accessibility profiles for these platelet progenitor cell types^55,92^. By partitioning platelet traits with GLEANR, we stratified these cell-type effects into factors which distinguish stages of platelet development. This demonstrates that GLEANR factors can support characterization of trait-relevant cell-type processes and development shared across GWAS phenotypes.

GLEANR has a few limitations. For example, enforcing sparsity in genetic factors may remove small effect or low-confidence signals, limiting its applicability to very rare traits, studies with small sample sizes, or those with a paucity of associations. In addition, our analysis does not account for every source of confounding between studies, such as assortative mating. However, if estimates of the impact of these confounders are available, GLEANR could be extended to adjust for those effects by integrating those estimates into 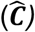. Finally, estimates of polygenicity and selection signatures from GLEANR factors are not numerically comparable to those from full GWAS, as they are based on the subset of variants chosen for inclusion in the analysis due to trait-association. Interpretation is thus limited to comparison with other factors in the same analysis.

In conclusion, GLEANR is a powerful tool for evaluation of pleiotropic components underlying genetic correlations. It can be applied to diverse GWAS and is available as a free software package along with supporting tools for interpretation of genetic factors. GLEANR enables characterization of trait genetic architectures and biological pathways into distinct components that clarify the pleiotropic impact of genetic associations across the human phenome.

## Acknowledgements

The authors would like to thank Battle lab members for helpful discussion throughout the course of this work. We especially thank Seraj Grimes and April Kim for manuscript feedback and code review, and Ben Strober for contributions to early ideation of project analyses. A.B. was supported by NIH/NIGMS award R35GM139580. M.A. was supported by NHLBI grant K08HL166690.

## Web resources

**GLEANR package:** https://github.com/aomdahl/gleanr

**Pan-UKBB:** https://pan.ukbb.broadinstitute.org

**Finngen:** https://finngen.gitbook.io/documentation/r10/releases

**Genetic Correlation Portal:** https://ukbb-rg.hail.is

**PheWeb summary statistics**: https://pheweb.org

**Neale Lab GWAS:** http://www.nealelab.is/uk-biobank/

**Tissue-specific S-LDSC Resources:** https://github.com/bulik/ldsc/wiki/Cell-type-specific-analyses

**Open Targets Portal:** https://genetics-docs.opentargets.org/data-access/data-download

## Author contributions

Ashton Omdahl: Conceptualization, Software, Methodology, Formal analysis, Data Curation, Writing-original draft, Writing – review & editing

Joshua Weinstock: Methodology, Writing – review & editing, Supervision

Rebecca Keener: Writing – review & editing, Project Administration

Surya Chhetri: Methodology

Marios Arvanitis: Conceptualization, Writing – review & editing, Supervision

Alexis Battle: Conceptualization, Resources, Writing – review & editing, Supervision

## Data and code availability

- Code pipelines used for analysis of published GWAS data and simulations are available at https://github.com/aomdahl/gleanr_workflow/.
- The GLEANR R package is available at https://github.com/aomdahl/gleanr.
- URLs for download of all FinnGen, PheWeb, Pan-UKBB and Neale Lab GWAS summary statistics used in this analysis are available at https://github.com/aomdahl/gleanr_workflow/.
- The genotype reference panel used for LD-pruning and clumping is available through the Thousand Genomes Portal: https://www.internationalgenome.org/data/.
- SNP-to-gene scores used in this study are available for download from Open Targets (Oct. 2022 release; Table S5): https://genetics-docs.opentargets.org/data-access/data-download.

