## Supplemental Notes, Figures, and Tables for "Sparse matrix factorization robust to sample sharing across GWAS reveals interpretable genetic components"

### Supplemental Information

#### Supplemental Note 1

##### GLEANR model fitting procedure

When fitting GLEANR, alternating regression steps are performed by fitting the following regression equations, implemented using sparse matrix functions in *glmnet*[1] in R 4.2.3:

Fitting U:

$$\mathrm{vec}(\boldsymbol{B}^*) = \widehat{\boldsymbol{V}}^* \times \mathrm{vec}(\boldsymbol{U}) + \epsilon, \qquad \text{subject to} \sum_{i,j=1}^{N,K} \left|U_{i,j}\right| < t_\alpha \tag{1}$$

Fitting V:

$$\mathrm{vec}(\boldsymbol{B}^*) = \widehat{\boldsymbol{U}}^* \times \mathrm{vec}(\boldsymbol{V}) + \epsilon, \qquad \text{subject to} \sum_{i=1,j=2}^{M,K} \left|V_{i,j}\right| < t_\lambda \tag{2}$$

Where:

- $\mathbf{B}^* = (\boldsymbol{W}_s \odot \mathbf{B})\boldsymbol{W}_c^T$ is an $M \times N$ matrix standardized to global zero mean, and unit variance.
- $\widehat{\boldsymbol{V}}^* = [(\boldsymbol{I_N} \otimes \boldsymbol{W_c})\boldsymbol{S_D^{-1}}] \otimes \widehat{\boldsymbol{V}}$, an $NM \times NK$ block diagonal matrix, with the $i^{th}$ diagonal block element corresponding to $\boldsymbol{W_c}\left(\mathrm{diag}(\boldsymbol{s_i})\right)^{-1}\widehat{\boldsymbol{V}}$.
- $\widehat{\boldsymbol{U}}^* = [(\boldsymbol{I_N} \otimes \boldsymbol{W_c})\boldsymbol{S_D^{-1}}]\left(\boldsymbol{I_M} \otimes \widehat{\boldsymbol{U}}\right)_N$, an $NM \times MK$ matrix. $\left(\boldsymbol{I_M} \otimes \widehat{\boldsymbol{U}}\right)_N \in \mathbb{R}^{NM \times KM}$ refers to the Kronecker product of the $M \times M$ identity matrix and $\widehat{\boldsymbol{U}}$, with rows re-ordered such that the top M rows correspond to rows containing $\boldsymbol{u_1^T} \in \mathbb{R}^{1 \times K}$ diagonally arranged, all the way down to $\boldsymbol{u_N}$, as in:

$$\begin{bmatrix} \boldsymbol{u_1^T} & \mathbf{0} & \mathbf{0} \\ \mathbf{0} & \boldsymbol{u_1^T} & \mathbf{0} \\ \mathbf{0} & \mathbf{0} & \ddots \\ \boldsymbol{u_2^T} & \mathbf{0} & \mathbf{0} \\ \mathbf{0} & \boldsymbol{u_2^T} & \mathbf{0} \\ \vdots & \vdots & \ddots \\ \boldsymbol{u_N^T} & \mathbf{0} & \mathbf{0} \\ \mathbf{0} & \boldsymbol{u_N^T} & \mathbf{0} \\ \mathbf{0} & \mathbf{0} & \ddots \end{bmatrix}$$

- $\boldsymbol{W}_s \in \mathbb{R}^{N \times M}$ contains weights corresponding to standard errors, $W_{s_{i,j}} = \frac{1}{s_{i,j}}$.
- $\boldsymbol{S_D^{-1}} \in \mathbb{R}^{NM \times NM}$, a block diagonal matrix, with the $i^{th}$ diagonal block corresponding to $\mathrm{diag}(\boldsymbol{s_i})^{-1}$.
- "vec" indicates stacking of matrix elements by row, here to yield a vector of size $NM \times 1$, and $\odot$ refers to the element-wise matrix product.
- $\boldsymbol{I_q}$ refers to the identity matrix of dimension $q \times q$.

To show that alternating fittings of Equations 1 and 2 fit the same global objective loss function, it is sufficient to show that $\boldsymbol{V}^* \times \mathrm{vec}(\boldsymbol{U}) = \boldsymbol{U}^* \times \mathrm{vec}(\boldsymbol{V})$:

$$\widehat{\boldsymbol{V}}^{*} \times \mathrm{vec}(\boldsymbol{U}) = \left[(\boldsymbol{I_N} \otimes \boldsymbol{W_c})\boldsymbol{S_D^{-1}}\right] \otimes \widehat{\boldsymbol{V}} \times \mathrm{vec}(\boldsymbol{U}) = \begin{bmatrix} \boldsymbol{W_c S_1^{-1} V} & \boldsymbol{0} & \boldsymbol{0} \\ \boldsymbol{0} & \boldsymbol{W_c S_2^{-1} V} & \boldsymbol{0} \\ \boldsymbol{0} & \boldsymbol{0} & \ddots \\ \vdots & \vdots & \vdots \end{bmatrix} \begin{bmatrix} \boldsymbol{u_1} \\ \boldsymbol{u_2} \\ \vdots \end{bmatrix}$$

$$= \begin{bmatrix} \boldsymbol{W_c S_1^{-1} V u_1} \\ \boldsymbol{W_c S_2^{-1} V u_2} \\ \vdots \end{bmatrix}$$

$$\widehat{\boldsymbol{U}}^{*} \times \mathrm{vec}(\boldsymbol{V}) = \left[(\boldsymbol{I_N} \otimes \boldsymbol{W_c})\boldsymbol{S_D^{-1}}\right]\left(\boldsymbol{I_M} \otimes \widehat{\boldsymbol{U}}\right)_N = \begin{bmatrix} \boldsymbol{W_c S_1^{-1}} & \boldsymbol{0} & \boldsymbol{0} \\ \boldsymbol{0} & \boldsymbol{W_c S_2^{-1}} & \boldsymbol{0} \\ \boldsymbol{0} & \boldsymbol{0} & \ddots \\ \vdots & \vdots & \vdots \end{bmatrix} \begin{bmatrix} \boldsymbol{u_1^T} & \boldsymbol{0} & \boldsymbol{0} \\ \boldsymbol{0} & \boldsymbol{u_1^T} & \boldsymbol{0} \\ \boldsymbol{0} & \boldsymbol{0} & \ddots \\ \boldsymbol{u_2^T} & \boldsymbol{0} & \boldsymbol{0} \\ \boldsymbol{0} & \boldsymbol{u_2^T} & \boldsymbol{0} \\ \vdots & \vdots & \ddots \end{bmatrix} \begin{bmatrix} \boldsymbol{v_1} \\ \boldsymbol{v_2} \\ \vdots \end{bmatrix}$$

$$= \begin{bmatrix} \boldsymbol{W_c S_1^{-1}} & \boldsymbol{0} & \boldsymbol{0} \\ \boldsymbol{0} & \boldsymbol{W_c S_2^{-1}} & \boldsymbol{0} \\ \boldsymbol{0} & \boldsymbol{0} & \ddots \\ \vdots & \vdots & \vdots \end{bmatrix} \begin{bmatrix} \boldsymbol{u_1^T v_1} \\ \boldsymbol{u_1^T v_2} \\ \vdots \\ \boldsymbol{u_2^T v_1} \\ \vdots \\ \boldsymbol{u_N^T v_M} \end{bmatrix} = \begin{bmatrix} \boldsymbol{W_c S_1^{-1}} & \boldsymbol{0} & \boldsymbol{0} \\ \boldsymbol{0} & \boldsymbol{W_c S_2^{-1}} & \boldsymbol{0} \\ \boldsymbol{0} & \boldsymbol{0} & \ddots \\ \vdots & \vdots & \vdots \end{bmatrix} \begin{bmatrix} \boldsymbol{V u_1} \\ \boldsymbol{V u_2} \\ \vdots \end{bmatrix}$$

$$= \begin{bmatrix} \boldsymbol{W_c S_1^{-1} V u_1} \\ \boldsymbol{W_c S_2^{-1} V u_2} \\ \vdots \end{bmatrix} \blacksquare$$

#### Updating the BIC to compare across settings of $K_{init}$

Calculation of BIC (Equations 5 and 6 in the main text) requires estimation of $\sigma^2$, which scales model error by the estimated residual variance of the data. GLEANR estimates $\sigma^2$ using the unbiased[2] ordinary least squares (OLS) residual variance estimator, in which OLS with the maximum number of input features under consideration ($NK$ for $\boldsymbol{U}$, $MK$ for $\boldsymbol{V}$) in a given iteration of GLEANR fitting is used to calculate residuals (as in the scikit-learn[3] 1.5.2 package). However, when comparing across settings of $K_{init}$, estimates of $\sigma^2_{OLS}$ may differ, as the number of free parameters grows linearly with $K_{init}$ (e.g. $NK_{init}$ for an estimate of $\boldsymbol{U}$). To account for this, when comparing $L_{BIC}$ across settings of $K_{init}$, GLEANR adopts the estimate of $\sigma^2_{OLS}$ corresponding to the greatest number of free parameters (typically for $K = M - 1$) for both $BIC_\alpha$ and $BIC_\lambda$, as all other models are "nested" within this one. GLEANR re-scales the fit terms for all $L_{BIC}$ before comparison across settings $K_{init}$

$$L_{BIC-global}(K) = BIC_{\lambda-global} + BIC_{\alpha-global} \tag{3}$$

$$= \frac{\left\|\mathbf{B}^{*} - \boldsymbol{U}^{*}\widehat{\boldsymbol{V}}^{\boldsymbol{T}}\right\|}{\sigma^2_{max-MK}} + \log(MN)\left\|\widehat{\boldsymbol{V_K}}\right\|_{0,0} + 2\gamma\log\begin{pmatrix} p \\ \left\|\widehat{\boldsymbol{V}}\right\|_{0,0}\end{pmatrix}$$
$$+ \frac{\left\|\mathbf{B}^{*} - \widehat{\boldsymbol{U}}\boldsymbol{V}^{*\boldsymbol{T}}\right\|}{\sigma^2_{max-NK}} + \log(MN)\left\|\widehat{\boldsymbol{U_K}}\right\|_{0,0} + 2\gamma\log\begin{pmatrix} p \\ \left\|\widehat{\boldsymbol{U}}\right\|_{0,0}\end{pmatrix} \tag{4}$$

where $\left\|\widehat{\boldsymbol{X}}\right\|_{0,0} = \sum_{n=1}^{N}\|\boldsymbol{x_n}\|_0$ is an unbiased estimate for the degrees of freedom of the lasso[4], and $\gamma = 1 - \log(NM)/2\log(P)$ is a scaling term that controls the contribution of the added BIC extension term[5]. $p$ is the total number of covariates under consideration ($NK$ for $\boldsymbol{U}$ or $MK$ for $\boldsymbol{V}$). We note that in some instances, the selected $\sigma^2_{max}$ is much smaller than expected, compared to estimates from other $K_{init}$ settings. This is likely due to overfitting of the data and results in the selection of high variance models. GLEANR is equipped to detect such cases and provides a user warning; users may wish manually review reports of $L_{BIC-global}$ scores to select more appropriate model settings.

### Supplemental Note 2

#### Matching factors to compare factorizations

When comparing matrix factorization outputs in both our simulation and cross-cohort comparative analyses, matrix factors (e.g. columns) are not guaranteed to be in the same order nor to be scale invariant. In order to compare output matrices, we matched matrix columns (factors) to each other by finding the factor order ($\boldsymbol{I_o}$) and sign vector ($\boldsymbol{\mu}$) that maximizes the global correlation across both matrices by optimizing the following:

$$\max_{\boldsymbol{I_o},\boldsymbol{\mu}}\left(\left(\mathrm{diag}\left[\mathrm{cor}\left(\boldsymbol{V},\boldsymbol{\mu}\boldsymbol{I_o}\widehat{\boldsymbol{V}}\right)\right]+\mathrm{diag}\left[\mathrm{cor}\left(\boldsymbol{U_1},\boldsymbol{\mu}\boldsymbol{I_o}\widehat{\boldsymbol{U}}\right)\right]\right)^2\right) \tag{5}$$

where $\boldsymbol{I_o}$ is an orthogonal rotation matrix with exactly one non-zero element per column of value 1, $\boldsymbol{\mu}$ is a vector of signs with elements $\mu_i \in [1,-1]$, and $\mathrm{cor}(\mathbf{X_1},\mathbf{X_2})$ is the $K \times K$ correlation matrix of the correlation between columns of $\mathbf{X_1} \in \mathbb{R}^{N\times K}$ and $\mathbf{X_2} \in \mathbb{R}^{N\times K}$. When the number of columns (i.e. $K$) of input matrices didn't match, columns were padded with zeros.

For matrices with small numbers of columns (e.g. $K < 8$), Equation 5 can be globally maximized by considering every possible permutation of column and sign order. However, for large $K$ this becomes combinatorially intractable ($2^K K!$ possible combinations). When $K > 6$, we used a greedy approach to maximize Equation 5 by first calculating a matrix $\mathbf{P} = (\mathrm{cor}(\boldsymbol{V_1},\boldsymbol{V_2}) + \mathrm{cor}(\boldsymbol{U_1},\boldsymbol{U_2}))^2$. We selected the entry of $\mathbf{P}$ with the maximal value ($\mathrm{P_{i,j}} = \max(\mathbf{P})$), and assigned factor $i$ in the target factorization (in $\boldsymbol{V}_1/\boldsymbol{U}_1$) to factor $j$ in the reference factorization (in $\boldsymbol{V}_2/\boldsymbol{U}_2$). We set column $j$ of $\mathbf{I_o}$ to a value of 1 in row $i$ with zeroes everywhere else, and $\mu_j = sign(\max(\mathrm{cor}(\boldsymbol{V}_1,\boldsymbol{V_2})_{i,j}, \mathrm{cor}(\boldsymbol{U_1},\boldsymbol{U_2})_{i,j})$. We then set all elements in row $i$ and column $j$ of $\mathbf{P}$ to zero and repeated this process with the next maximal element of $\mathbf{P}$. We repeatedly matched factors in this manner until only six columns/factors remain, at which point we solve for the remaining columns of $\boldsymbol{I_o}$ and entries of $\boldsymbol{\mu}$ by globally maximizing Equation 5.

### Supplemental Note 3

#### Trait selection for GLEANR evaluation of 137 diverse traits

From 7228 published phenotypes evaluated by the Pan-UKBB[6], we filtered down to just 137 for inclusion in our analysis. We first removed traits failing certain Pan-UKBB consortium quality control standards, including a valid LDSR[7] $h^2 \in (0,1)$ with a significant Z-score, normal $\lambda_{gc}$ across top 3 powered ancestries, and an LDSR inflation ratio < 0.3. We further filtered to traits with $N_{eff} > 5000$ (where $N_{eff}$ is the harmonic mean of cases and controls for categorical traits and the total number of samples for continuous traits) and those with LDSR liability scale $h^2 > 0.05$. This left 736 GWAS phenotypes, including several in categories which would be challenging to interpret and not of interest in this analysis, which we excluded. These groups included activities (e.g. transportation preferences, phone side use), environment (e.g. type of work or income), nutrition (e.g. bread preference, coffee preference), social interactions (e.g. leisure activities), operation codes, consultant treatment specialties, hospital admission codes, parental ages, prescriptions, acquired phenotypes (e.g. acquired toe deformities), and some psychiatric traits (e.g. alcohol with meals, cannabis use). Category assignments were based on previously reported domain assignments[8], extended to include UKBB coding information (Table S2). We also excluded studies performed on raw continuous phenotypes (those not estimated using inverse rank normal transformed values), leaving 474 studies. To avoid including phenotypes with redundant sources of genetic variation in the human phenome, we further filtered remaining phenotypes by their pairwise genetic correlation. Estimates of genetic correlation came from XT-LDSR[9] as downloaded from the Neale Lab Genetic Correlation Browser[10] (Web Resources). In order of decreasing heritability, we added traits to our set which had an estimated $r_g < 0.7$ with every other trait already in our set, yielding 165 phenotypes. These we supplemented with 26 biomarker and 22 other phenotypes which did not have published $r_g$ estimates available (including hyperlipidemia, pulse pressure, indirect bilirubin, low density lipoprotein). After eliminating phenotypes with redundant mappings between the Pan-UKBB and the Genetic Correlation Browser (preferring the study with the largest $N_{eff}$), we performed XT-LDSC across HapMap3 SNPs on all pairs of 194 remaining GWAS. Beginning with the phenotype with the largest estimate of heritability and proceeding in descending order of liability scale heritability, we added GWAS to our final set for analysis set such that no two traits had pairwise $r_g < 0.7$, yielding 137 traits.

#### LD-independent variant selection for evaluation of 137 diverse traits

To select approximately LD-independent variants for inclusion in our analysis, we performing clumping using plink[11] on a set of 309,191 HapMap3 SNPs with MAF > 0.01. Typically when clumping, variants within a given LD block are prioritized for inclusion by a pre-specified heuristic, such as the p-value of association in a GWAS study. To favor inclusion of variants more likely to harbor multi-trait pleiotropic signal, we used a score of pleiotropy for variants as the variant prioritization heuristic. This was calculated for SNP $n$ as:$h_n = 1 - (\frac{1}{N+1}\sum_{m=1}^{M} \mathrm{I}_{[p_{n,m}<1\times10^{-5}]})$, where $\mathrm{I}_{[p_{n,m}<1\times10^{-5}]}$ is an indicator function returning 1 if the GWAS p-value for SNP $n$ of trait $m$ passes below our nominal threshold of $p < 1 \times 10^{-5}$. Variants achieving this threshold in more studies would have smaller $h_n$ and so be prioritized by the clumping procedure.

#### Supplemental figures:

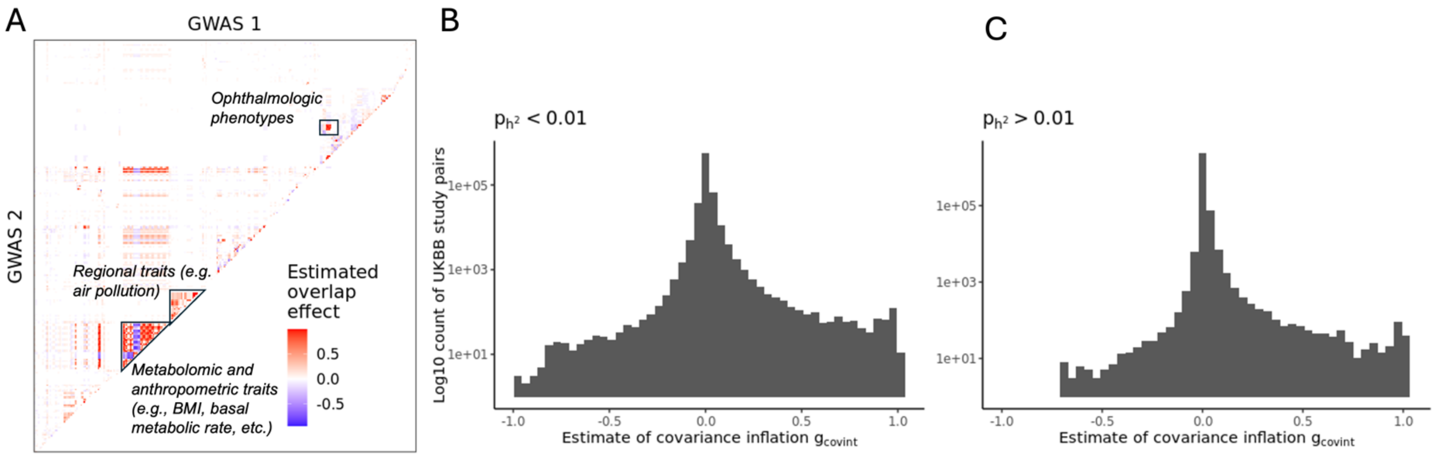


**Figure S1: Estimates of cohort overlap between GWAS evaluated in the UKBB.** A) Heatmap of cross-trait LD score regression[9] estimates of covariance inflation due to sample sharing ($g_{covint}$) from the Neale Lab Genetic Correlation browser[10] across 292 traits included in the 2019 Tanigawa et al.[12] study, matched using fuzzy string matching of phenotype names. Several prominent block structures exist, including commonly evaluated anthropometric phenotypes. Only trait pairs with $p < 0.05$ Bonferroni-corrected significant estimates are visualized. B) Histogram of $g_{covint}$ estimates across all pairwise GWAS tested in the Neale Lab Genetic Correlation browser download[10] with valid heritability estimates at $p < 0.01$ (N=700,550 pairs of GWAS phenotypes). Counts are given on the log scale. Many pairs of heritable traits have evidence of covariance inflation due to sample sharing. C) Histogram of $g_{covint}$ estimates across all pairwise tested studies in the Neale Lab Genetic Correlation browser download[10] which had valid heritability estimates with $p > 0.01$ (N=2,473,335 pairs of GWAS phenotypes), with counts given on the log scale.

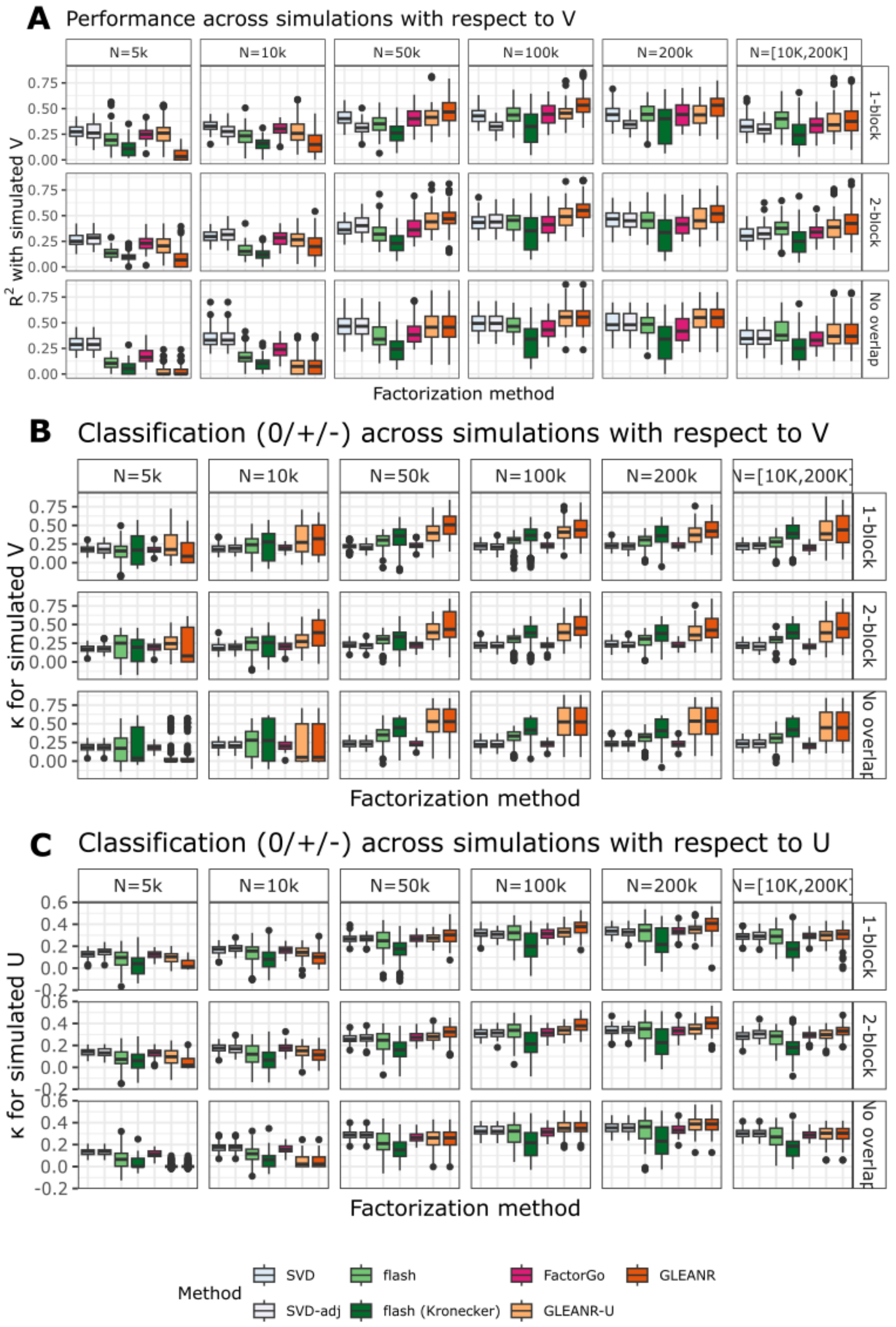


**Figure S2: Additional metrics of performance of GWAS matrix factorization methods on simulated GWAS summary statistics.** A) Boxplot of factor variance explained ($R^2$) between estimated $\hat{\boldsymbol{V}}$ matrices and simulated $\boldsymbol{V}$. Simulation results correspond to those for $\boldsymbol{U}$ in Figure 2 of the main text. B and C) Classification similarity of factors between estimated and simulated $\boldsymbol{V}$ (in B) and $\boldsymbol{U}$ (in C), across tested matrix factorization methods. Similarity was measured using Cohen's Kappa ($\kappa$), with matrix elements assigned to one of three classes as either positive, negative, or zero. Because of the sign variance across factorizations, the highest scoring classification for positive and negatively assigned matrix elements were considered.

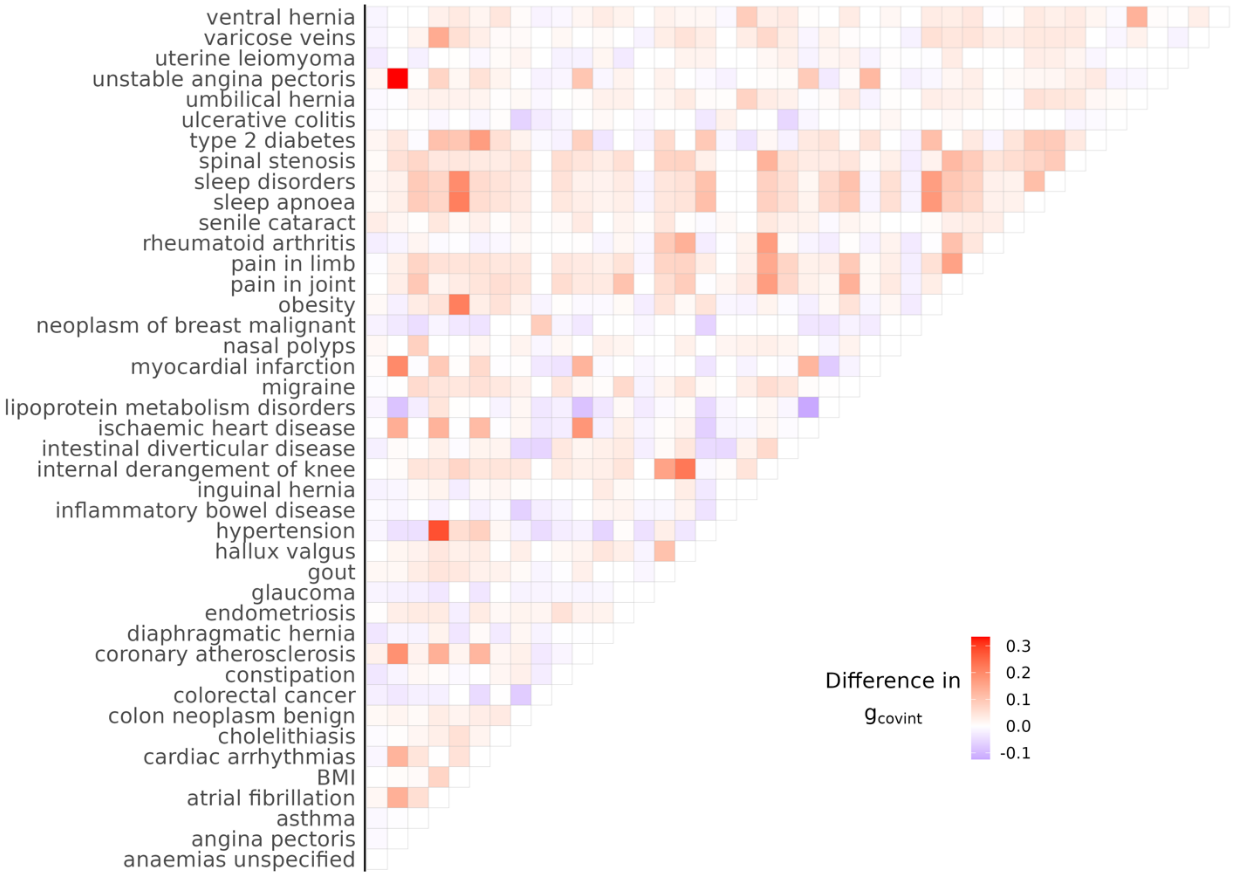


**Figure S3: Heatmap of differences in raw XT-LDSC estimates of covariance inflation due to sample sharing ($g_{covint}$) across all pairs of 42 traits evaluated in both FinnGen and UKBB GWAS**. Heatmap reports $g_{FinnGen} - g_{UKBB}$.

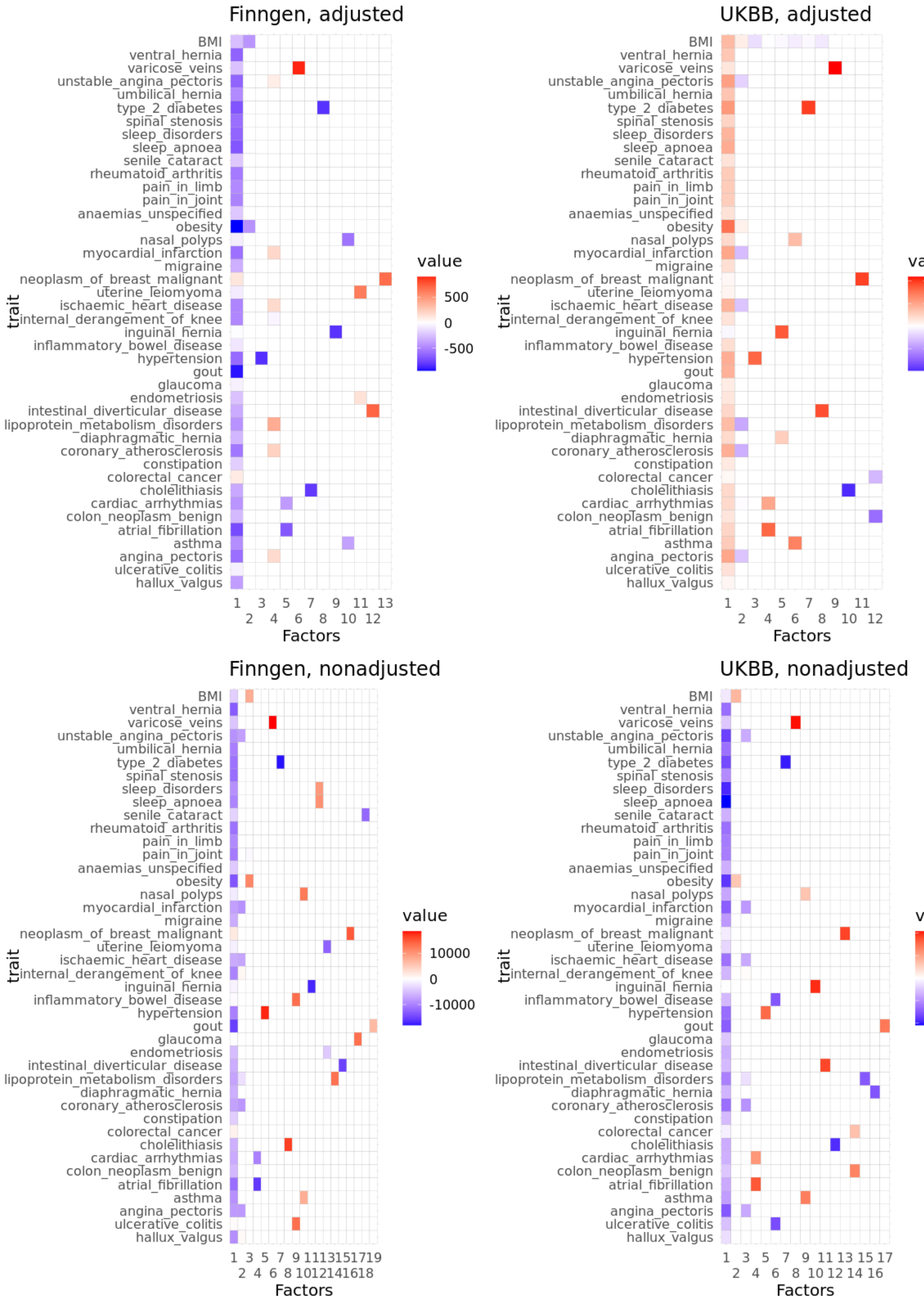


**Figure S4: Factorizations of 42 traits by GLEANR from FinnGen and UKBB GWAS, with and without covariance adjustment.** Top heatmaps were estimated with GLEANR, and lower two were estimated with GLEANR-U. Heatmaps on the left came from FinnGen, and those on the right form UKBB GWAS. Color indicates the contribution of a given phenotype to a particular factor.

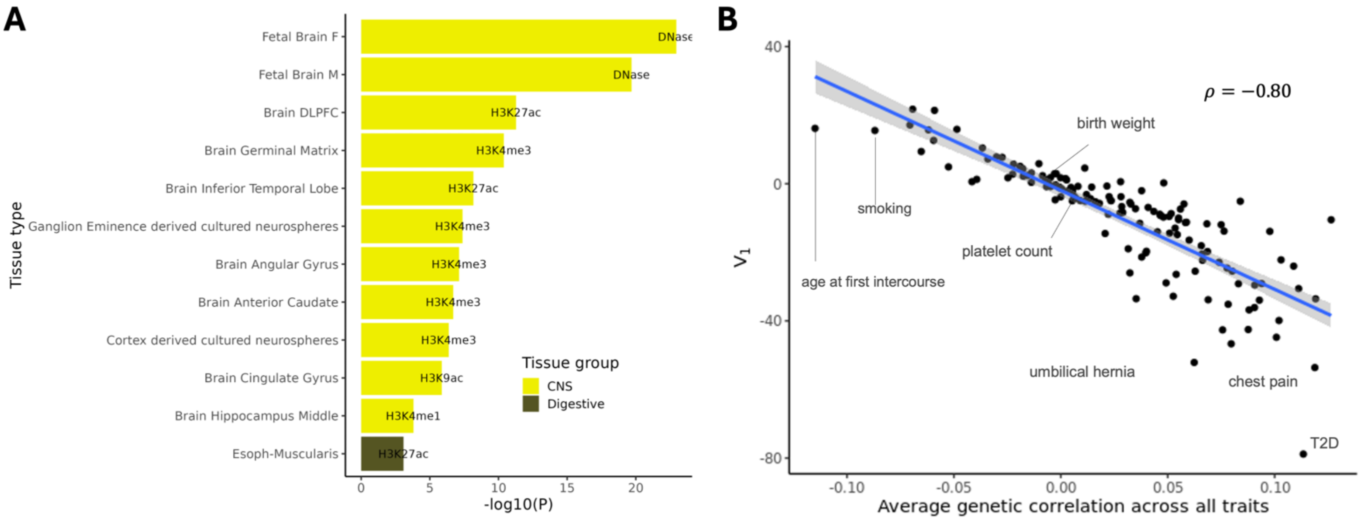


**Figure S5: Factor 1 was enriched for tissue-specific chromatin markers and correlated strongly with the average genetic correlation per trait.** A) Top 12 tissue-specific chromatin marker enrichments from S-LDSR[13] for Factor 1. Label on bar indicates the type of marker tested, bar length corresponds to the -log10 p-value, and color indicates to the tissue group of interest. B) Average estimates of $r_g$ by XT-LDSR for all 137 input traits (x-axis), plotted against $V_1$ weight (y-axis). Average $r_g$ per trait was calculated as the mean of all $r_g$ estimates with FDR > 0.05 corresponding to that trait. Reported $\rho = -0.80$ indicates the overall correlation between raw values of $V_1$ and and average $r_g$ per trait.

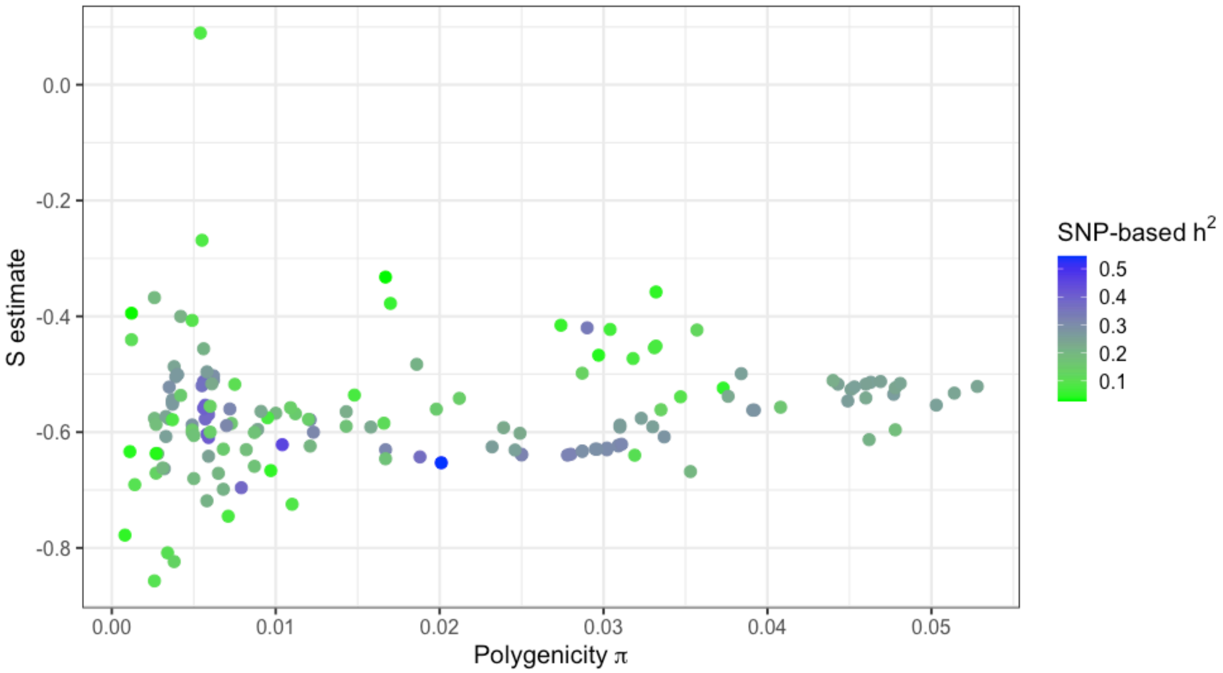


**Figure S6: Estimates of polygenicity ($\pi$, x-axis), signature of selection ($S$,y-axis) and SNP-based heritability (point color) from SBayesS for 154 GWAS phenotypes as reported by Zeng et al, 2021[14].** Traits with the largest magnitude $S$ tend to have reduced polygenicity estimates, while those with large polygenicity estimates have low or medium $S$. A number of traits with low polygenicity also reflect weak signatures of negative selection.

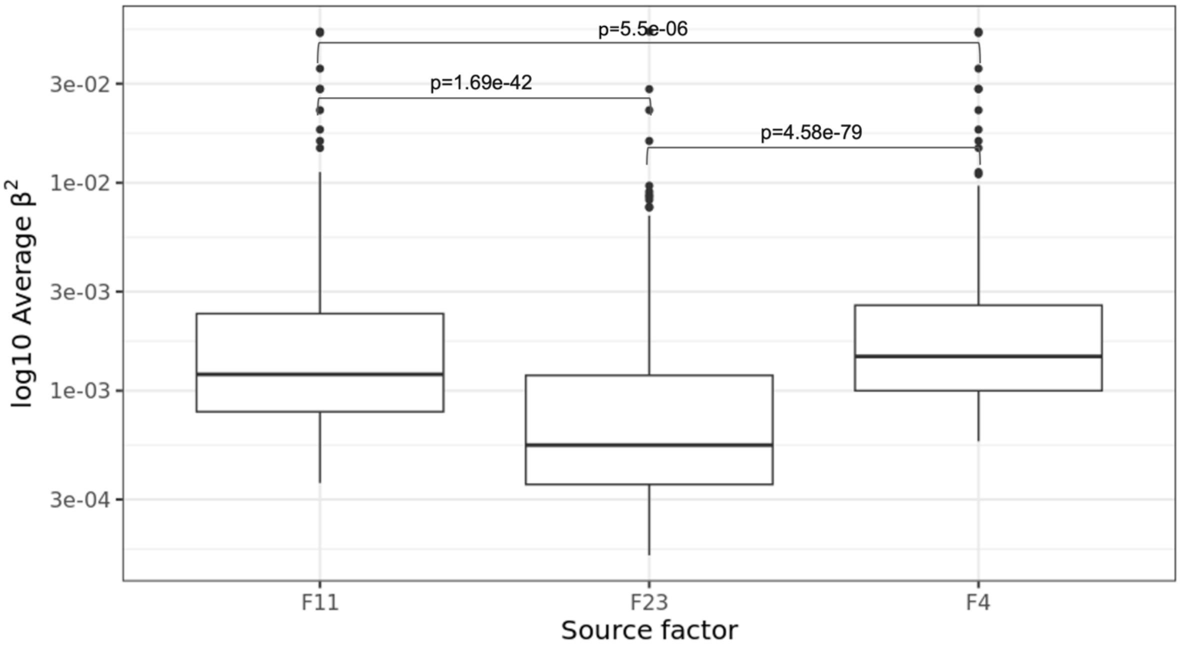


**Figure S7: Distribution of average size magnitudes for top SNPs in Factors 4, and 11 and 23 across mean platelet volume, platelet count, and platelet distribution width GWAS.** Boxplots reflect the log10 average GWAS $\beta^2$ for top factor SNPs from each factor. Top F4 SNPs have larger effect size magnitudes per allele on average than both Factors 11 and 23. Effect size estimates used in this visualization were those from European Pan-UKBB[6] GWAS used in our analysis of 137 diverse traits. P-values come from a pairwise Wilcoxon rank sum test, Benjamini-Hochberg corrected for multiple tests.

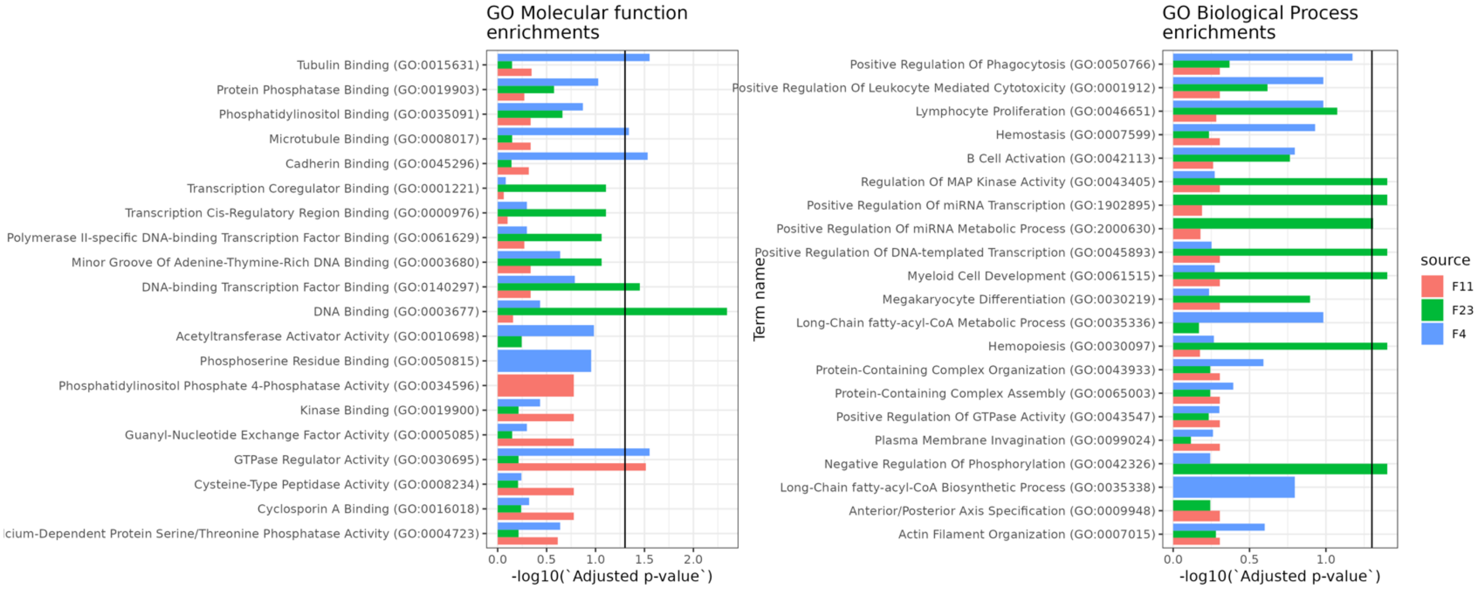


**Figure S8: Top GO molecular function and biological process term enrichments reported for top factor genes in Factors 4,11, and 23.** Bars indicate the -log10 Benjamini-Hochberg adjusted p-value for enrichment, and colors correspond to source factor. Black line indicates an adjusted p-value of 0.05. GO terms selected for visualization are any appearing in the top seven most strongly enriched for any factor. Note that some enrichment estimates were not available for some factors.

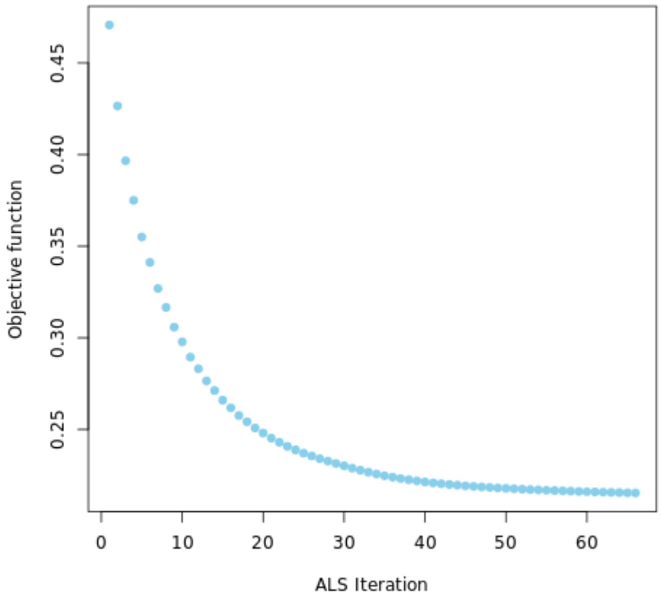


**Figure S9: Objective function at each alternating least squares iteration during model fitting of GLEANR.** Fitting visualized here came from analysis of 137 diverse UKBB phenotypes reported in the text. A single alternative least squares (ALS) iteration (x-axis) refers to an updated fitting of either $U$ or $V$.

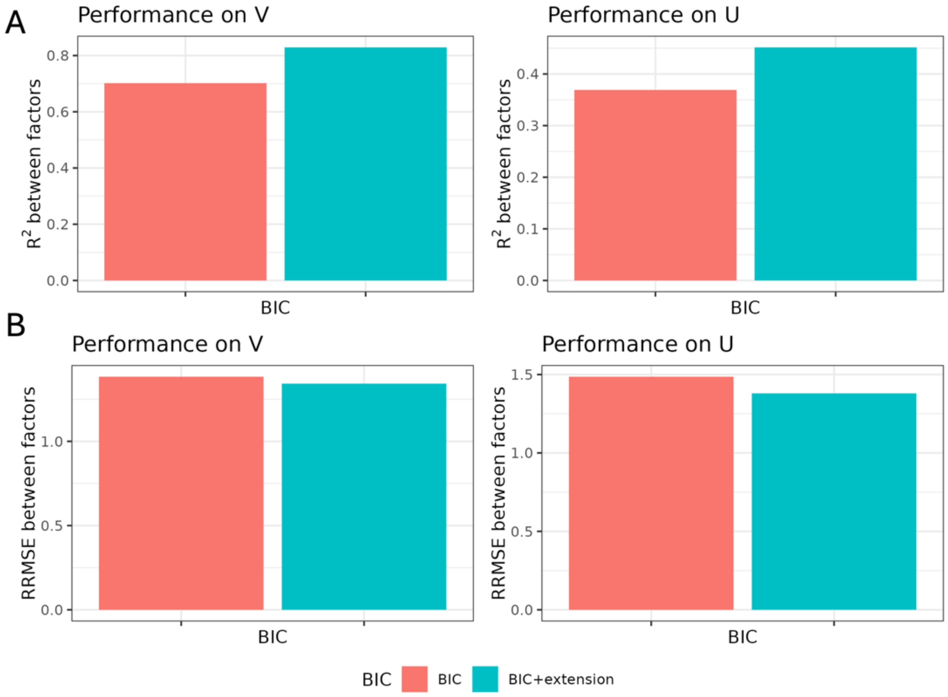


**Figure S10: Comparison of reproducibility of GLEANR factors across cohorts with different implementations of the BIC (with and without the extension adjusting for large parameter spaces).** Plots indicate metrics of the comparison of factors estimated from GWAS in UKBB or FinnGen. A) Bars indicate the $R^2$ between factors in either cohort. In both $\boldsymbol{U}$ and $\boldsymbol{V}$, BIC with the extension adjusting for large sample spaces[5] yields more similar factors across the UKBB and Finngen. B) Bars indicate the relative root mean squared error. For both $\boldsymbol{U}$ and $\boldsymbol{V}$, BIC with the extension results in a lower error.

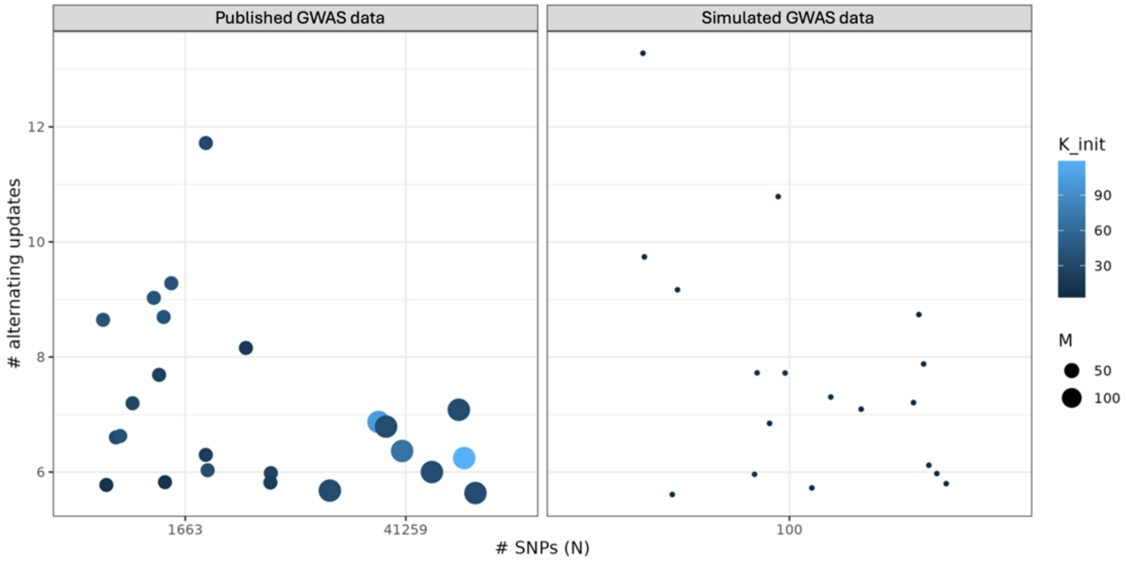


**Figure S11: Dotplot of number of GLEANR model selection iterations to achieve convergence of $L_{BIC}$ with respect to $M, N$ and $K_{init}$.** One dot represents a single instance performing model selection with GLEANR. Dot position corresponds to the number of SNPs evaluated ($N$, x-axis) and the total number of alternating iterations before achieving convergence (y-axis, minimum five). Dot size corresponds to the number of traits under evaluation ($M$), while color corresponds to the number of factors $K_{init}$ chosen for initialization. Overall, most model selection procedures converge within ten alternating iterations, regardless of sample size.

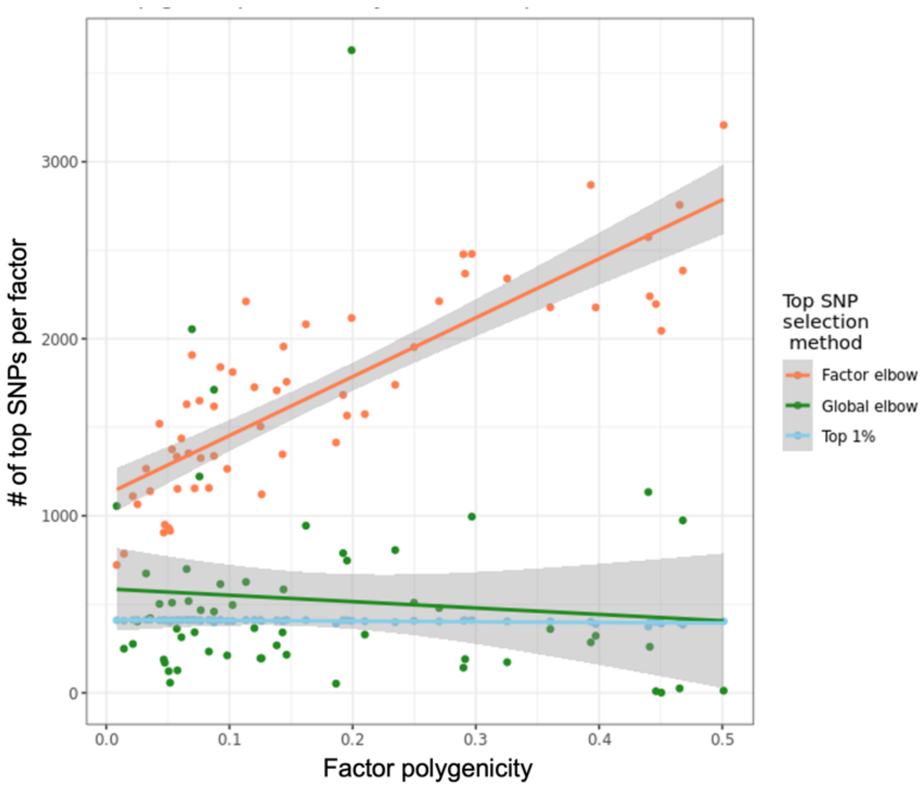


**Figure S12: Relationship between factor polygenicity estimate (x-axis) and the number of top factor SNPs nominated by different prioritization methods (y-axis).** Nomination of top SNPs by the global elbow (green) refers to selecting an effect size threshold for elements of $\boldsymbol{U}$ across all factors that corresponds to the “elbow” point of a scree plot of ranked absolute weights in $\boldsymbol{U}$. The “Top 1%” method designates genes corresponding to just the top 1% of SNPs with the largest magnitude in every factor. The factor elbow method (red) corresponds to selecting SNPs which appear above the “elbow” point of a scree plot of ranked absolute weights for each factor individually.

| Factor | SNP | chr | pos | Factor rank | Open Targets[15,16] Gene | GWAS evidence | Regulatory evidence |
|---|---|---|---|---|---|---|---|
| **F4** | rs2289420 | 3 | 124667095 | 1 | KALRN | Mean platelet volume, platelet count, platelet distribution width, gamma glutamyl transferase levels | |
| | rs2228367 | 19 | 16086510 | 2 | TPM4 | Linked to lead GWAS variants in mean platelet volume, platelet count, platelet distribution width, platelet crit | Blood eQTL in eQTLGen (p=6.9e-91) and GTEx whole blood (p=1.9e-7) for TPM4 |
| | rs16843769 | 1 | 172062471 | 3 | DNM3 | Platelet count | VEP intronic variant; blood eQTL for METTL13 in eQTLGen (p=2.0e-7) |
| **F11** | rs6092755 | 20 | 59248274 | 1 | ZNF831 | Nominal associations in reticulocyte count (p = 2.2e-5) and platelet count (3.9e-5) | Blood eQTL for ZNF831 in eQTLGen (p=1.1e-30) |
| | rs16843769 | 1 | 172062471 | 2 | DNM3 | Platelet count | VEP intronic variant; blood eQTL for METTL13 in eQTLgen (p=2.0e-7) |
| | rs6673413 | 1 | 171927810 | 3 | METTL3 | Platelet count | Blood eQTL for METTL3 in eQTLGen (p=4.1e-25) |
| **F23** | rs2239196 | 12 | 111435363 | 1 | ALDH2 | White blood cell count, platelet count, eosinophil count, eosinophil percent, reticulocyte count, | Blood eQTL for ALDH2 in eQTLGen (p=1.7e=69) |
| | rs10744774 | 12 | 111652218 | 2 | ALDH2 | White blood cell count, eosinophil percent of granulocytes, eosinophil percent of WBCs, hemoglobin concentration, red blood cell count | Blood eQTL for ALDH2 (p=1.7e=69) and TMEM116 (3.0e-30) in eQTLGen |
| | rs2228367 | 19 | 16,086,510 | 3 | TPM4 | Linked to lead GWAS variants in mean platelet volume, platelet count, platelet distribution width, platelet crit | Blood eQTL in eQTLGen (p=6.9e-91) and GTEx whole blood (1.9e-7) for TPM4 |

**Table S8: Regulatory evidence supporting top SNPs in platelet-related factors (Factors 4, 11, and 23).** SNPs were mapped to genes using Open Targets[15,16] weighted ranking integrating multiple lines of evidence. Cited GWAS studies are those listed with a $p < 5 \times 10^{-8}$ on the Open Targets online portal; if there were more than five, only the top five most significant are listed. Regulatory evidence includes eQTLs reported on Open Targets.

| Platelet Development Stage | Disease category | Source | Genes |
|---|---|---|---|
| All | Hereditary macrothrombocytopenia (HMTP) | Collins et al, 2021. "Advances in understanding the pathogenesis of hereditary macrothrombocytopenia." British Journal of Haematology | THPO, ANKRD26, ETV6, FLI1, GATA1, GFI1B, HOXA11, MECOM, RUNX1, NBEAL2, ABCG5, ABCG8, GNE, MPIG6B, SLFN14, SRC, ACTB, ACTN1, CDC42, DIAPH1, FLNA, MYH9, TUBB1, GP1BA, GP1BB, GP9, ITGA2B, ITGB3, VWF |
| Early megakaryopoiesis | Inherited platelet disorders | Lentaigne et al, 2016. "Inherited platelet disorders: toward DNA-based diagnosis." Blood. | THPO, MPL, GATA1, RUNX1, FLI1, ETV6, GFI1B, HOXA11, MECOM, ANKRD26, RBM8A |
| Late megakaryopoiesis | Inherited platelet disorders | Collins et al, 2021. "Advances in understanding the pathogenesis of hereditary macrothrombocytopenia." British Journal of Haematology; Lentaigne et al, 2016. "Inherited platelet disorders: toward DNA-based diagnosis." Blood. | P3B1, HPS1, HPS3, HPS4, HPS5, HPS6, BLOC1S3, BLOC1S6, DTNBP1, LYST, VPS33B, VIPAS39, STXBP2, NBEA, NBEAL2, CYCS, SRC, SLFN14, PLAU, STIM1,ABCG5, ABCG8, GNE, MPIG6B |
| Proplatelet formation | Inherited platelet disorders | Collins et al, 2021. "Advances in understanding the pathogenesis of hereditary macrothrombocytopenia." British Journal of Haematology; Lentaigne et al, 2016. "Inherited platelet disorders: toward DNA-based diagnosis." Blood. | MYH9, WAS, ACTN1, FLNA, TUBB1, DIAPH1, GP1BA, GP1BB, GP9, ITGA2B, ITGB3, VWF, ACTB, CDC42 |
| Platelet activity | Inherited platelet disorders | Lentaigne et al, 2016. "Inherited platelet disorders: toward DNA-based diagnosis." Blood. | P2RY12,TBXA2R, TBXAS1, PLA2G4A, ITGA2B, RASGRP2,VWF, ITGB3, FERMTS, GP1BA,GP9, GP1BB, GP6, ANO6 |

**Table S9: Genes related to inherited platelet disorders (including hereditary macrothrombocytopenia) that correspond to stages of platelet formation**. Reported genes were used in enrichment tests of genes from Factors 4,11, and 23. Gene lists and stage of platelet development were drawn from the two publications as listed (“Source” column).

| Proportion shared, cohort 1 | Proportion shared, cohort 2 | Proportion shared, cohort 3 | Overall proportion shared (out of 90,000) |
|---|---|---|---|
| 0.000 | 0.000 | 0.000 | 0.000 |
| 0.333 | 0.000 | 0.000 | 0.111 |
| 0.500 | 0.000 | 0.000 | 0.167 |
| 0.667 | 0.000 | 0.000 | 0.222 |
| 0.333 | 0.333 | 0.000 | 0.222 |
| 0.767 | 0.000 | 0.000 | 0.256 |
| 0.500 | 0.333 | 0.000 | 0.278 |
| 0.867 | 0.000 | 0.000 | 0.289 |
| 1.000 | 0.000 | 0.000 | 0.333 |
| 0.667 | 0.333 | 0.000 | 0.333 |
| 0.500 | 0.500 | 0.000 | 0.333 |
| 0.333 | 0.333 | 0.333 | 0.333 |
| 0.767 | 0.333 | 0.000 | 0.367 |
| 0.667 | 0.500 | 0.000 | 0.389 |
| 0.500 | 0.333 | 0.333 | 0.389 |
| 0.867 | 0.333 | 0.000 | 0.400 |
| 0.767 | 0.500 | 0.000 | 0.422 |
| 1.000 | 0.333 | 0.000 | 0.444 |
| 0.667 | 0.667 | 0.000 | 0.444 |
| 0.667 | 0.333 | 0.333 | 0.444 |
| 0.500 | 0.500 | 0.333 | 0.444 |
| 0.867 | 0.500 | 0.000 | 0.456 |
| 0.767 | 0.667 | 0.000 | 0.478 |
| 0.767 | 0.333 | 0.333 | 0.478 |
| 1.000 | 0.500 | 0.000 | 0.500 |
| 0.667 | 0.500 | 0.333 | 0.500 |
| 0.500 | 0.500 | 0.500 | 0.500 |
| 0.867 | 0.667 | 0.000 | 0.511 |
| 0.767 | 0.767 | 0.000 | 0.511 |
| 0.867 | 0.333 | 0.333 | 0.511 |
| 0.767 | 0.500 | 0.333 | 0.533 |
| 0.867 | 0.767 | 0.000 | 0.544 |
| 1.000 | 0.667 | 0.000 | 0.556 |
| 1.000 | 0.333 | 0.333 | 0.556 |
| 0.667 | 0.667 | 0.333 | 0.556 |
| 0.667 | 0.500 | 0.500 | 0.556 |
| 0.867 | 0.500 | 0.333 | 0.567 |
| 0.867 | 0.867 | 0.000 | 0.578 |
| 1.000 | 0.767 | 0.000 | 0.589 |
| 0.767 | 0.667 | 0.333 | 0.589 |
| 0.767 | 0.500 | 0.500 | 0.589 |
| 1.000 | 0.500 | 0.333 | 0.611 |
| 0.667 | 0.667 | 0.500 | 0.611 |
| 1.000 | 0.867 | 0.000 | 0.622 |

| | | | |
|---|---|---|---|
| 0.867 | 0.667 | 0.333 | 0.622 |
| 0.767 | 0.767 | 0.333 | 0.622 |
| 0.867 | 0.500 | 0.500 | 0.622 |
| 0.767 | 0.667 | 0.500 | 0.644 |
| 0.867 | 0.767 | 0.333 | 0.656 |
| 1.000 | 1.000 | 0.000 | 0.667 |
| 1.000 | 0.667 | 0.333 | 0.667 |
| 1.000 | 0.500 | 0.500 | 0.667 |
| 0.667 | 0.667 | 0.667 | 0.667 |
| 0.867 | 0.667 | 0.500 | 0.678 |
| 0.767 | 0.767 | 0.500 | 0.678 |
| 0.867 | 0.867 | 0.333 | 0.689 |
| 1.000 | 0.767 | 0.333 | 0.700 |
| 0.767 | 0.667 | 0.667 | 0.700 |
| 0.867 | 0.767 | 0.500 | 0.711 |
| 1.000 | 0.667 | 0.500 | 0.722 |
| 1.000 | 0.867 | 0.333 | 0.733 |
| 0.867 | 0.667 | 0.667 | 0.733 |
| 0.767 | 0.767 | 0.667 | 0.733 |
| 0.867 | 0.867 | 0.500 | 0.744 |
| 1.000 | 0.767 | 0.500 | 0.756 |
| 0.867 | 0.767 | 0.667 | 0.767 |
| 0.767 | 0.767 | 0.767 | 0.767 |
| 1.000 | 1.000 | 0.333 | 0.778 |
| 1.000 | 0.667 | 0.667 | 0.778 |
| 1.000 | 0.867 | 0.500 | 0.789 |
| 0.867 | 0.867 | 0.667 | 0.800 |
| 0.867 | 0.767 | 0.767 | 0.800 |
| 1.000 | 0.767 | 0.667 | 0.811 |
| 1.000 | 1.000 | 0.500 | 0.833 |
| 0.867 | 0.867 | 0.767 | 0.833 |
| 1.000 | 0.867 | 0.667 | 0.844 |
| 1.000 | 0.767 | 0.767 | 0.844 |
| 0.867 | 0.867 | 0.867 | 0.867 |
| 1.000 | 0.867 | 0.767 | 0.878 |
| 1.000 | 1.000 | 0.667 | 0.889 |
| 1.000 | 0.867 | 0.867 | 0.911 |
| 1.000 | 1.000 | 0.767 | 0.922 |
| 1.000 | 1.000 | 0.867 | 0.956 |
| 1.000 | 1.000 | 1.000 | 1.000 |

**Table S13: Proportion of samples shared when estimating GWAS summary statistics from each of the three simulated cohorts in simulation scenario one**. Right-most column ("Overall proportion shared (out of 90,000)") indicates what proportion of the total number of possible shared samples were used to estimate GWAS (max possible 90,000), calculated as the sum of re-used samples in all cohorts.
